# Cortical ductility governs cell-cell adhesion mechanics

**DOI:** 10.1101/2023.11.28.568975

**Authors:** Aditya Arora, Mohd Suhail Rizvi, Gianluca Grenci, Florian Dilasser, Chaoyu Fu, Modhura Ganguly, Sree Vaishnavi, Kathirvel Paramsivam, Srikanth Budnar, Ivar Noordstra, Alpha S. Yap, Virgile Viasnoff

**Affiliations:** Mechanobiology Institute, National University of Singapore, 5a Engineering drive 1, 117411 Singapore; Department of Biomedical Engineering, Indian Institute of Technology Hyderabad, Telangana, India; Institute for Molecular Bioscience, The University of Queensland, St. Lucia, Brisbane, Queensland, Australia 4072; CNRS, IRL3639, 5a Engineering drive 1, 117411 Singapore; CSL Innovation Pty Limited at the Bio21 Molecular Science and Biotechnology Institute, The University of Melbourne, Parkville, Victoria 3010, Australia

## Abstract

This paper challenges our understanding of cell-cell adhesion by emphasising the role of mechanical dissipation at the cellular level. We have developed new microdevices to measure the energy dissipated during the rupture of junctions between cell-cell doublets. Using a synthetic cadherin approach, we decoupled the role of cadherin binding energy, signalling and downstream regulation of cytoskeletal architecture. This yielded a phase diagram in which cell junctions transition from a ductile to a brittle fracture mode based on their ratio of cortical tension and shape relaxation time. We recapitulated our results using a descriptive mechanical simulation approach. Our results shift our understanding of cell-cell adhesion from the current focus on bond energy and tension to the key role played by energy dissipation in the cytoskeleton during junction deformation and its active mechanosensitive regulation.

## Introduction

Epithelial tissues are constantly exposed to internal and external forces both during development, and under homeostasis ^1, 2^. The maintenance of physical integrity and barrier function despite these stresses is central to the physiological function of epithelia ^3^. Cell-cell adhesions such as the adherens junction, which mechanically couples the actomyosin cytoskeleton of cells through series of adaptor proteins, serve to maintain tissue integrity while transmitting force over long distances ^4^. While much is known about adherens junction formation and its interaction with the cytoskeletal network, our understanding of how cells resist mechanically disruptive stress is still incomplete. A better understanding of the basis and regulation of tissue cohesion is crucial to explain its role in processes such as morphogenesis and diseases such as cancer.

The paradigmatic understanding of cell-cell adhesion stability is described by the free energy cost of establishing a junction. It is usually described as the sum of the adhesion energy due to the binding affinity ^5, 6^, and to the mechanical tension in the cortex ^7, 8^ that is accounted as an effective elastic energy ^5, 6, 8, 9^. This effective elastic energy is a descriptive proxy that is not explicitly related to any measured biochemical properties, and is usually serves as a fitting parameter to successfully predict the static conformation of the cells in the tissue. However, de-adhesion or detachment processes occur at different rates, and often under non-equilibrium conditions and therefore cannot be explained by such an equilibrium force balance. During cell detachment, cells undergo notable deformation, this process involves both elastic and dissipated energy, part of which is liberated after de-adhesion when cells rapidly round up ^9^. To study the role of the rheological properties of cortex in regulating the resistance of the cell-cell contacts to mechanical rupture, it is important to realise that this problem has almost exclusively been studied from the perspective of the maximum force (called adhesion strength)^9–11^ that the junction can withstand. The interpretation of these measurements are underpinned by the view that mechanical resistance is due to the molecular bonds formed by E-cadherin either in *trans* ^6^ or with the cortex ^8^. This bond-centered approach is convenient; however, it has always been impossible to relate the microscopic forces measured to break cadherin bonds to the maximum force a junction can withstand.

In soft matter physics, fracture resistance is known to depend on a more complex set of material parameters. These include viscoelasticity and plasticity in addition to binding energy and elasticity. Material toughness, i.e. the total energy required to fracture the material, is proving to be a more appropriate descriptor of a material’s ability to resist mechanical fracture. To our knowledge, the toughness of a cell-cell contact has never been determined, so its influence on cell detachment is often ignored. However, it is now well understood how large dissipative energies render certain hydrogels fracture resistant ^12^. It has been shown that optimal polymer network properties allow energy to be dissipated from an adhesion interface or crack into the bulk of a hydrogel material. This can support extremely strong bonds, independent of the bond strength per se. ^12, 13^. Thus, energy dissipation not only acts as an additional rate-dependent viscous component, but may also redistribute mechanical stress to stabilise the integrity of the adherens zone. We hypothesised that, analogous to dissipative polymeric hydrogels, the isotropic fibrous acto-myosin cortex attached to cadherin adhesions may act as an energy sink during the process of cell separation and thus play a deterministic role in controlling cell adhesion strength.

## Results

### Decoupling actin cortex properties from binder recruitment

Understanding the role of energy dissipation in actomyosin cortex is complicated by the strong intertwining of cadherin recruitment with cortex properties. ^14–16^. For example, cadherin recruitment leads to biochemical and biophysical modulation of cortical properties via various Rho GTPases ^16–18^. In turn, cortical tension drives the mechanosensitive recruitment of cadherins at adherens junction ^15^. To decouple the adhesive properties of the cadherins from their signaling, we first constructed chimeric cadherins by replacing the 5 EC domains in the extracellular region of of E-cadherin with a halo-tag and by mutating a conserved pro-endocytic di-leucine motif in the cytosolic domain ^19^. The transmembrane and the cytosolic domains were otherwise unchanged (see **Material and Methods**). Cadherin-null A431D cells were stably transfected with GFP-labelled version of this chimeric cadherin. Separate batches were incubated with complementary ssDNA tagged with Halo-ligand (**Figure 1a and b**) (**Material and Methods**). To minimize steric effects, the total length of the binder was kept constant and only the length of base-pairing region was varied. The chimeric DNA-cadherin (DNA-cad) junctions were >95% selective (**Figure 1c**) for forming junctions with complementary DNA strands.

**Fig 1.**
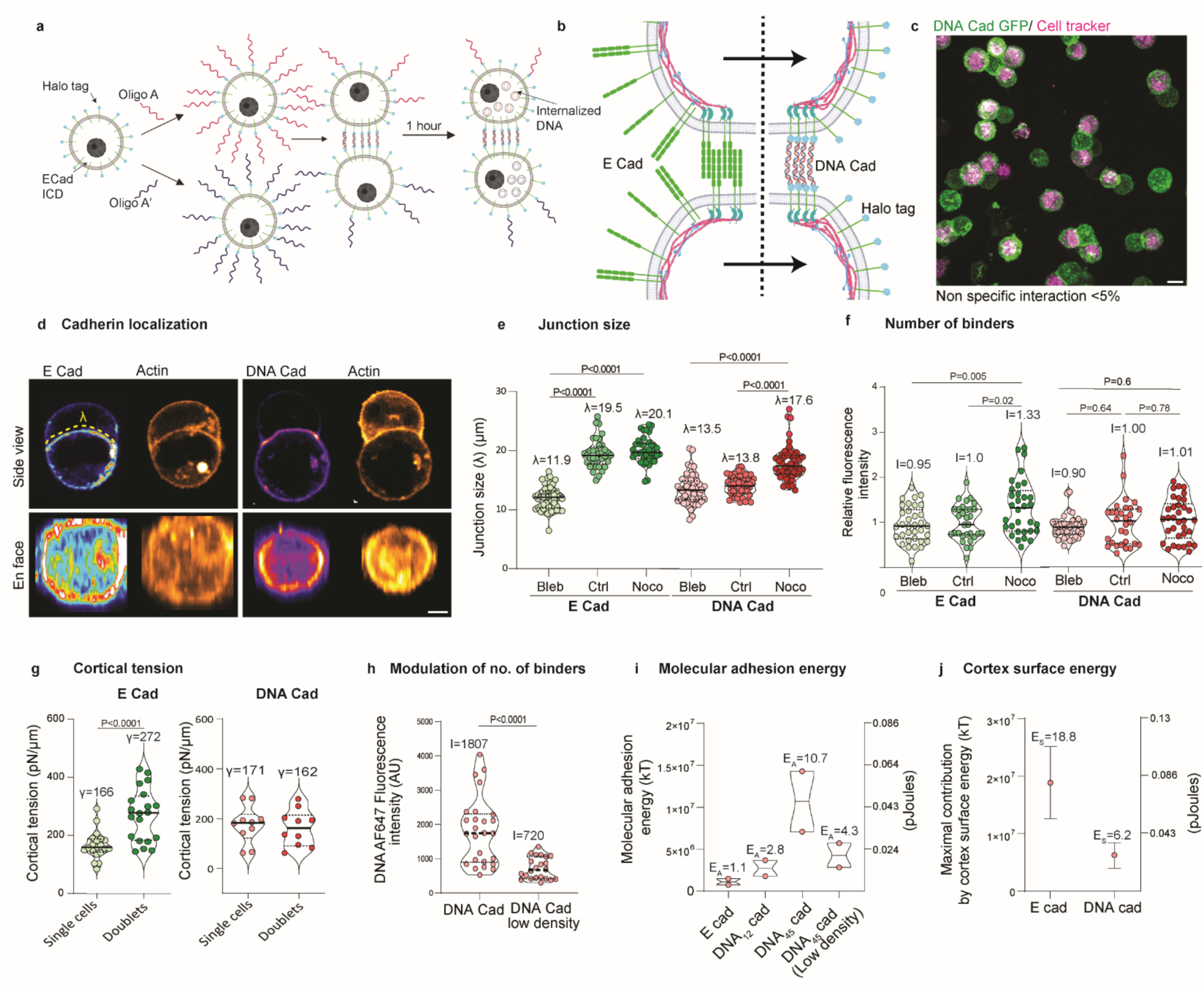
DNA-cadherin forms bonafide cell-cell adhesion. (a) Schematic of cell-cell adhesion formation based on DNA-cadherins wherein the ECD of Ecad is replaced with halo tag and used to covalently bind complementary ssDNA oligonucleotides. (b) schematic of cell-cell junction with either E-cad or DNA-cad, showing a key difference in terms of availability of free binder for stimulus dependent recruitment of new binder in case of E-cad. (c) Cells labelled with complementary DNA strands labelled with DNA-cad GFP alone and DNA-cad GFP with cell tracker, demonstrating contact formation between cells with complementary strands. (d) Side view and *en face* view of contact for cell doublets formed through E-cad (E cad GFP, and Lifeact mCherry); and DNA-cad (DNA – AF647, and Lifeact mCherry) under increased cortical tension using nocodazole (10 μM) treatment. Scale 5 μm. (e) Quantification of junction size for cell-cell doublets formed through E-cad and DNA-cad under blebbistatin (25 μM), control and nocodazole (10 μM) treatment. (f) Relative fluorescence intensity of E-cad GFP and DNA AF647 at the cell-cell contact under blebbistatin (25 μM), control and nocodazole (10 μM) treatment. (g) Cortical tension of single cells and cell doublets interacting through E-cad and DNA-cad. (h) DNA AF647 intensity after modulating number of ssDNA on cells using halo ligand ssDNA: halo ligand ratio of 3:1. (h) Comparison of molecular adhesion energies of E-cad and DNA-cad with 12 bp, 45 bp strands and 45 bp strands at reduced density. (i) Maximal energetic contribution due to the cortical tension at the contact surface.

We compared the morphologies of the adherens junctions formed between cell doublets expressing either WT E-cadherin or DNA-cad that were grown in suspension (**Figure 1d, Supplementary** Figure 1). In particular, the following features were identical for both cadherin systems: *i-* both binders were distributed to the edge of the cell contacts (**Figure 1d, and Supplementary** Figure 1) (N=15), *ii-* the tension dependent depletion of the actin cortex in the centre of the contact (**Supplementary figure 1**)(N=15) *iii*-the tension dependent increase of junction size (**Figure 1e**) *iv-* the dynamics of junction expansion with a characteristic time τ between 5 to 7 minutes (Supplementary Figure 2a) (N=15). It resulted that the organizations of binders and cytoskeleton were identical when we compared WT E-cad and chimeric cadherin coupled with any of the DNA lengths we used: 12bp (DNA*_12_*-cad), 20bp (DNA*_20_*-cad) and 45bp (DNA*_45_*-cad).

However, the recruitment of DNA-cad to the junction proved insensitive (P=0.78) (**Figure 1f**) to the modulation of myosin activity by either blebbistatin or nocodazole. Notably, whereas WT E-cad increased by 33 (± 11%; P=0.02) under increased mechanical tension, this did not occur with DNA-cad (**Figure 1f**). In fact, DNA-cad A431D has a finite pool of DNA binders that are either endocytosed or permanently bound at the junction 60 minutes after the junction elongation started (Supplementary Figure1). It showed that the DNA-cad junctions have a partial loss of mechanosensitive regulation of junctional components and maintain a constant total number of binders independent of the mechanical stimulation.

On the other hand, E-cad is also known to regulate cortical properties through outside-in signalling. This was confirmed by a significant upregulation of cortical tension after doublet formation in E-cad cells. (**Figure 1g**). In contrast, doublet formation in the presence of DNA-cad did not lead to any change in cortical tension, indicating a lack of biophysical regulation of the cortex after replacement of E-cad ectodomains (**Figure 1g**). Overall, DNA-cad adhesion produces junctions that are structurally bona fide adherens junctions with controllable adhesion energy (number of binders and energy per binder). However, they were decoupled from mechanosensitive binder turnover and essential outside-in signalling. (**Supplementary Table 1**). This provided the opportunity to use DNA-cad to precisely control adhesion energy by either i) using different lengths of DNA molecules or ii) modulating the density of DNA molecules on the cell surface by competitive labelling between ssDNA halo ligand and halo ligand alone (**Figure 1h**). Using these approaches, we found that cell-cell adhesion could be modulated over a wide range of molecular adhesion energies, i.e. from the order of E-cad adhesion to 10-fold higher, while holding other factors constant (**Figure 1i**). In addition to the molecular adhesion energy, we estimated the maximum energetic contribution from surface energy reduction at the contact to be significantly higher for E-cad vs. DNA-cad doublets (**Figure 1j**). Overall, DNA-cad provides a controlled platform to independently modify cortical properties and binder affinities while maintaining physiological relevance for understanding cell-cell adhesion mechanics.

### Measuring toughness of adherens junctions

We then used this experimental system to study the cohesivity of adherens junctions. We developed (**Materials and Methods**) a comb of cantilevers (**Figure 2a** **and Supplementary** Figure 3) to perform microstrain tensile tests mounted on an inverted microscope (**Figure 2a**). Cell doublets are attached at one end to the flexible cantilever, and the second cell is pulled at a constant rate (≈ 20μm/s) using a suction micropipette (**Figure 2b**). This allowed us to simultaneously measure the force (cantilever deflection) and deformation (doublet elongation) that the cell doublets undergo during detachment (**Figure 2b**). We calculated the separation force (Fs), the strain to failure (ε) and the toughness of the junction, i.e. the work done or the total energy under the curve required to separate the contact (U) (**Figure 2c**). The strain rate was set so that the junction breaks within 10 s to test the role of cytoskeletal architecture before major actin turnover can occur.

**Fig 2.**
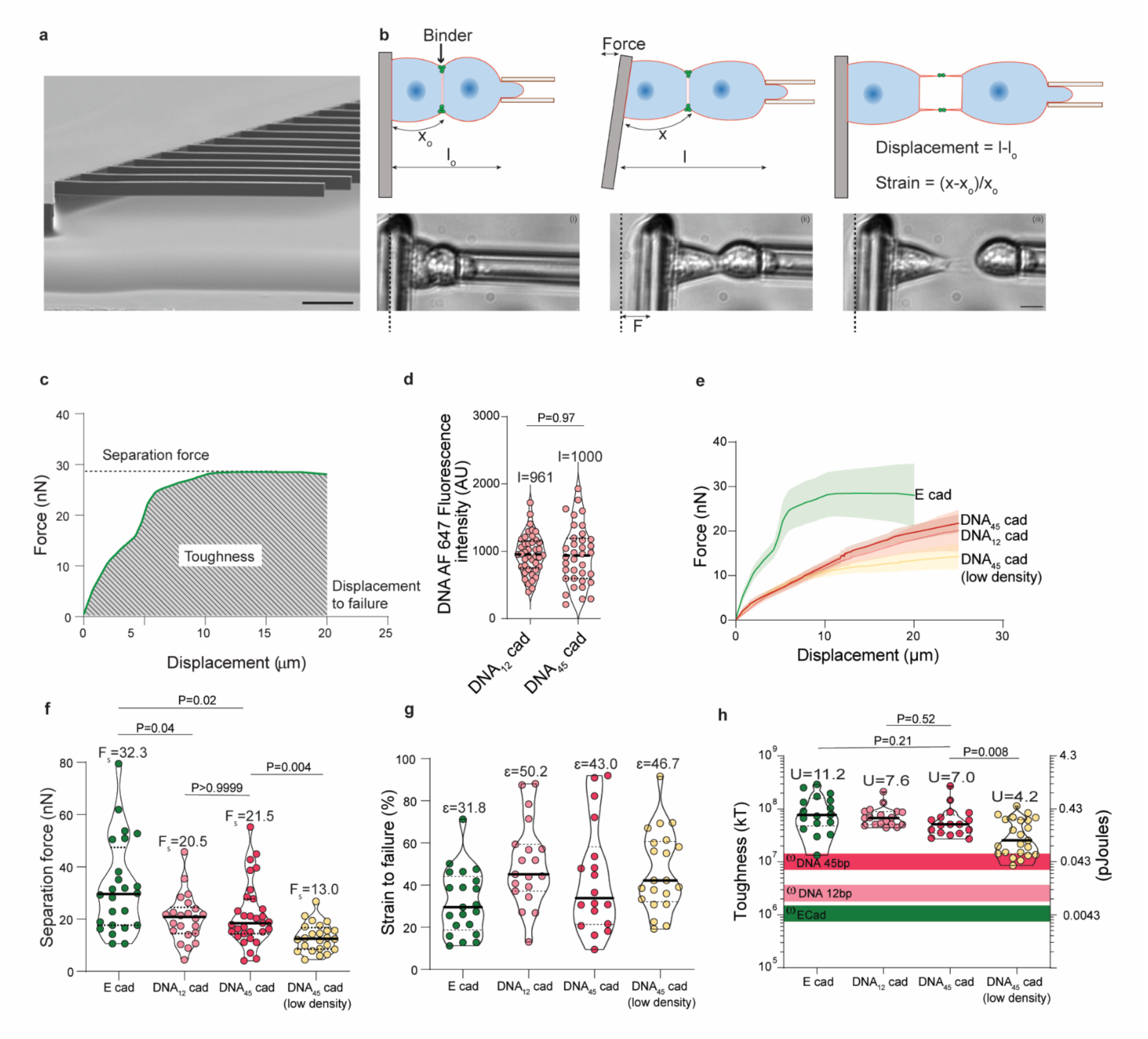
Cortex modulation and ligand number but not adhesive energy dictate cell adhesion strength. (a) Scanning electron micrograph of an SU-8 cantilever microchip array. Scale 200 μm. (b) Schematic of the different stages during cell separation experiments indicating force (F), displacement and strain (ε) measurements during the experiment, along with correlative examples of a cell doublet. Scale 10 μm. (c) Example of a force-displacement curve for a cell doublet depicting separation force (Fs) and toughness (U) of cell adhesion. (d) DNA AF647 fluorescence intensity at contacts formed by 12 and 45 bp DNA. (e) Averaged force displacement curves for E-cad, DNA_12_ cad, DNA_45_ cad and DNA_45_ cad low density cell doublets (shaded region represents 95% confidence interval of respective curves). (f) Separation force, (g) Strain to failure and (h) Toughness for E-cad, DNA_12_ cad, DNA_45_ cad and DNA_45_ cad low density cell doublets, the central line in violin plots represents the median and values indicated above each group represents the mean. The shaded horizontal bars in (h) indicate the calculated molecular adhesion energies (Ꙍ) for E-cad, DNA_12_ Cad, and DNA_45_ Cad.

### Binding energies of individual receptor hardly influence cell-cell adhesion

We first varied the *trans* binding free energies of individual cadherin binders while keeping the total number of binders constant (**Figure 2d**). We compared junctions formed with WT E-cad of 5-11kT/mol ^9^, 12bp DNA-Cad: 22kT/mol, and 45bp DNA-Cad: 85kT/mol ^20^. Based on the quantification of the total number of DNA-cad in the cells we estimated the total energy of the contact of the order of 1.1 x 10^6^, 2.8 x 10^6^, and 10.7 x 10^6^ kT for E-cad, DNA_12_ cad and DNA_45_ cad respectively (Figure 1i).

The force displacement curve showed the presence of a prominent plastic/ductile zone in the case of WT E-cad cell doublets (**Figure 2e**). The reference WT E-cad expressing A431D displayed a separation force F_s_ of 32 ± 17 nN, a rupture strain ε of 32 ± 15 % and Toughness U of 1.1 x 10^8^ kT (**Figure 2f-h**). For DNA-cad cells Fs dropped to 20 ± 9 nN (12bp) and 22 ± 12 nN (45bp), ε increased to 50% and 43% respectively. U remained similar for all cell types. Remarkably, the changes in cohesion of the junction were not dependent on the energy of the individual binders. In fact, the dissociation force was 50% higher for E-cad than for DNA-cad, despite its much lower binding affinity. Furthermore, all three parameters remained constant irrespective of the binding energy of the DNA strands between DNA12 and DNA45 cads. We therefore concluded that the binding energy per binder does not play a consequential role in the cohesion of the junction in this system. Previously, this was thought to be due to the small contribution of adhesion tension to the interfacial tension of the cell-cell contact compared to the cortical tension of the cells ^8^. However increasing the adhesive tension by even one order of magnitude in this case had no effect on adhesion toughness, indicating that adhesive energy plays a negligible role in cell detachment.

We then asked whether the total number of receptors influences cell adhesion strength. Reducing the number of DNA-cad molecules at the contact by 50% (**Figure 1h**) led to a significant decrease in both separation force (40%, P=0.01) and adhesion toughness (40%, P=0.003) (**Figure 2f and 2h**). Although the net molecular adhesion energy of DNA*_45_*-cad-low density group was higher than DNA*_12_*-cad (Figure 1i), both separation force and adhesion toughness were significantly higher for DNA*_12_*-cad than DNA*_45_*-cad-low density group (P<0.05). Therefore, the decrease in the adhesion strength for low density DNA_45_ cad is likely to be a result of the lower number of receptors rather than net adhesion energy. Overall, our data suggest that the binding strength of individual cadherin molecules plays a minor role in cell-cell adhesion strength. Hence, some other factor must be more critical.

### From ductile to brittle rupture of junctions

As an alternative to binding strength, it has been proposed that junctional stability results from cortical tension, which stabilises cadherin at the junction and leads to mechanosensitive actin recruitment. Since DNA-cad lacks mechanosensitive binder recruitment, we were able to test the role of cortical tension on cell doublet cohesion independently of stress-induced recruitment and signalling of trans-binding proteins. We used blebbistatin, calyculin A and nocodazole (**Material and Methods**) to modulat the cortical tension of DNA*_45_*-cad expressing cells by 15 fold from about 100 to 1500 pN/μm (**Figure 3a, Supplementary figure 4a**). As cortical tension increased, cell separation evolved from ductile (large deformation at constant stress) to brittle (abrupt rupture in the elastic regime) with a marked reduction in the plastic plateau in the stress-strain curve. (**Figure 3b** **– g, Supplementary videos 1-3**). The changes in the mode of fracture resulted in F_s_ peaking at a maximal separation force (31 ± 13 nN) at moderate cortical tension (700 pN/μm, Calyculin 100nM) (**Figure 3d**). Increasing tension further induced a fourfold drop down to 8 ± 4 nN (1500 pN/μm, nocodazole 10 μM). Further, true separation stress (separation force normalized by the junction size) never increased significantly but showed a drastic drop in the highest cortical tension group (**Supplementary figure 4b**). In contrast, the strain to failure ε anti-correlated with the cortical tension. Indeed blebbistatin-treated cells (ϒ=94 ± 38 pN/μm) showed a fourfold higher strain (66 ± 37 %) to failure compared to the highest concentration of nocodazole (16 ± 11%) (**Figure 3e**). As a result, the toughness of junctions plateaued for cortical tension up to around 700 pN/μm. It then drastically dropped from around 10^8^ kT to 6 x 10^6^ kT for the highest tension group. (**Figure 3f**). Thus, increasing tension compromised the ability of junctions to deform in response to stress and, indeed, promoted a catastrophic mode of brittle junction failure.

**Fig 3.**
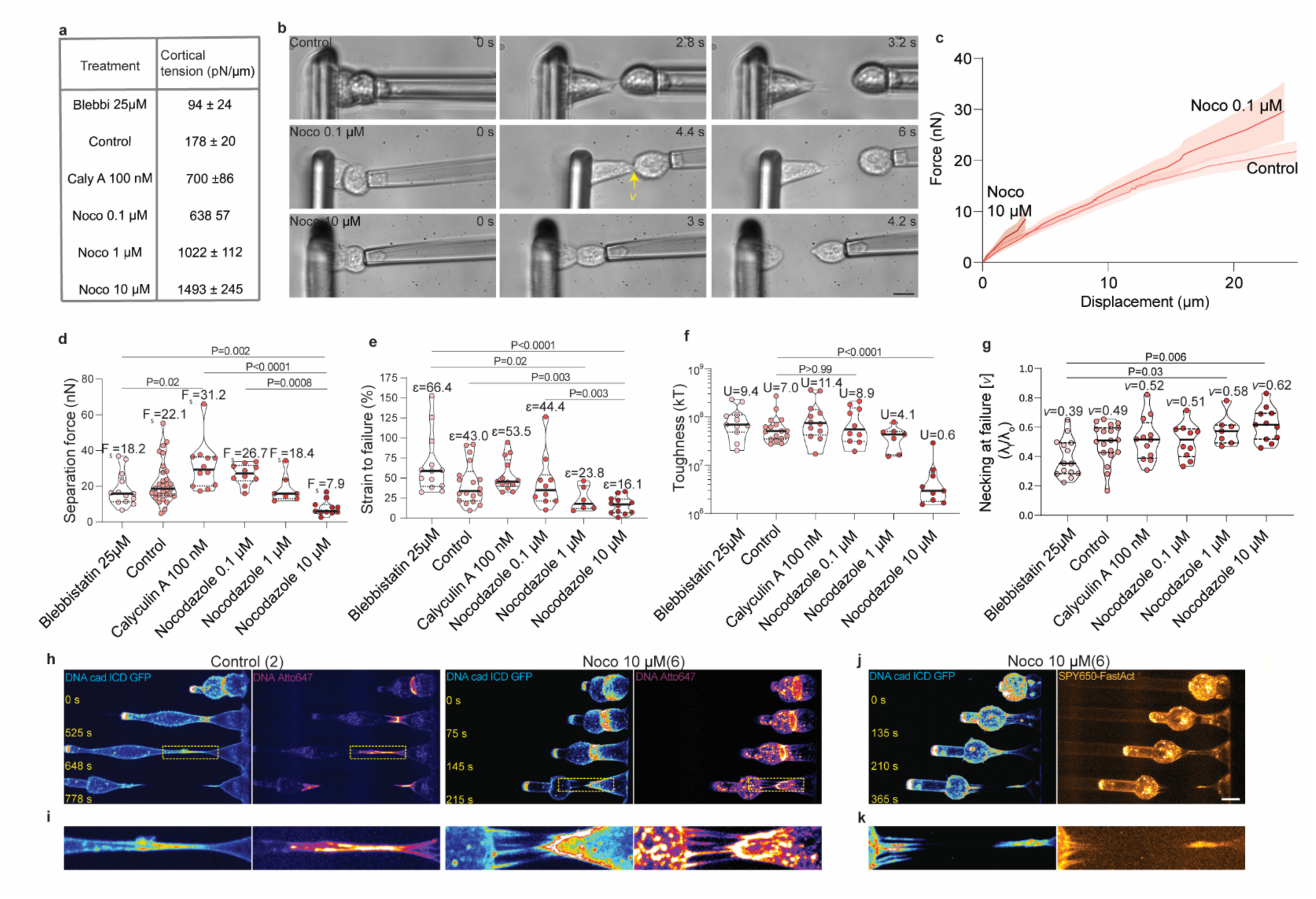
Contractility driven increase in cortical tension correlates positively with brittleness. (a) Cortical tension of DNA-cad expressing cells can be modulated over a broad range upon different drug treatments (values represented at mean ± SD. (b) Snapshots of DNA-cad doublets under different treatments during cell separation experiments. Scale 10 μm. (c) Averaged force displacement curves control, nocodazole 0.1 μM and 10 μM treated DNA-cad cell doublets (shaded region represents 95% confidence interval of respective curves). (d) Separation force, (e) Strain to failure, (f) Toughness and (g) extent of necking for DNA-cad doublets with increasing cortical tension, the central line in violin plots represents the median and values indicated above each group represents the mean. (h) Maximum intensity projections of Cad GFP and DNA Alexa fluor 647 signal from cells with low and highest tension values being pulled to cause junction fracture. Scale 10 μm. (i) Zoomed in view of the demarcated region in (h) at fracture. (j) Maximum intensity projections of Cad GFP and SPY650 FastAct signal from cells with highest tension value being pulled to cause brittle junction fracture. Scale 10 μm. (k) Zoomed in view of the demarcated region in (j) at fracture, demonstrates pulling out of actin filaments from cortex into the tethers.

We imaged the junctions using the GFP-tagged intracellular domains of the DNA-cad (labelling all the intracellular domains of the synthetic cadherin regardless of their coupling to a DNA strand), and using Alexa 647 fluorescent ssDNA as the DNA linkers for only one cell of the doublet. At low tension, the cortex proved deformable allowing the binders to funnel and concentrate into a shrinking contact which eventually failed with a single large tether (**Figure 3h and i**, **Supplementary video 4**). At higher tension, the cell cortex hardly deformed, the junction barely shrunk (**Figure 3g**) and a catastrophic fracture occurred as multiple tethers originated from the edges of the contact (**Figure 3h and i**, **Supplementary videos 5)**. In order to assess whether these tethers were caused by the detachment of DNA-cad from the cortex or by the failure of the actin network and the partial pulling out of actin filaments from the network, we imaged actin in the same experiment as above. It was clear that although the tethers did not have a homogeneous presence of actin along their length, they did show an enrichment of actin signal at their base and partial pulling of some actin filaments from the cortex into the tethers. (**Figure 2j and k, Supplementary video 6**). This suggested that a partial failure of the actin filament network in the cortex occurred during the junction fracture.

Overall, an increase in contractility in DNA*_45_*-cad cells led to a non-monotonic cohesion of the cell doublets. This suggests that the regulation of the energy dissipation during the plastic deformation of the cell cortex and/or cytoskeleton may be a potent mechanism by which cells can maintain high resistance to mechanical stress.

### Actin architecture governs cortical dissipative character, ductility and hence, adhesion strength

To test the hypothesis that the architecture of the actin cortex allows dissipation and plays a critical role in maintaining cell cohesion, we altered actin polymerisation and branching in DNA_45_-cad cells while limiting the junctional pool of binders. This allowed us to investigate how different actin nucleators affect cell-cell adhesion strength. To test the role of branched and linear actin polymerisation, cells were transfected with constitutively active (CA) versions of N-WASP (CA N-WASP) and mDia1 (CA mDia1). In both cases, the static size of the junction remained similar, with a small increase of less than 10% when the two nucleators were combined. (**Supplementary** Figure 5a and 5c). A significantly lower binder concentration was achieved for Cad GFP-CA-N-WASP fusion transfected cell lines, as higher expression levels proved toxic for making stable lines (**Supplementary figure 5b**).

As demonstrated by **Figure 4a** **(Supplementary videos 7)**, CA N-WASP fusion at the C terminus of DNA*_45_*-cad resulted in cells with highly deformable cortices with an almost two-fold increase in strain before failure (from 43% for control to 85%) (**Figure 4b and 4d**). Despite cells deforming into highly elongated cones that remained adherent at their common summit, they displayed no significant alteration of their separation force or toughness (**Figure 4c-e**) as compared to DNA*_45_*-cad only. By contrast, CA mDia1 co-expression with DNA*_45_*-cad led to a less substantial increase in cortex deformation (**Figure 4a, 4b and 4d, Supplementary video 8**) with a 50% increase in strain to failure (from 43% to 67%) accompanied by a large increase in separation force of around two-fold (from 21nN for Ctrl to 39nN for CA mDia1), and consequently a doubling of the adhesion toughness (from 7.10^7^ to 19.10^7^ kT). A double transfection with both CA N-WASP and CA mDia1 (**Material and Methods**) led to a synergistic effect on cortex properties reinforcing both its ductility (strain to failure reaching 94%), as well as separation force (40.8nN) hence exceeding the toughness (28.8 .10^7^ kT) of the control wild type E-cad junctions by two-fold (28.8.10^7^ kT *vs* 11.2. 10^7^ kT) (**Figure 4e and 2h, Supplementary video 9**).

**Fig 4.**
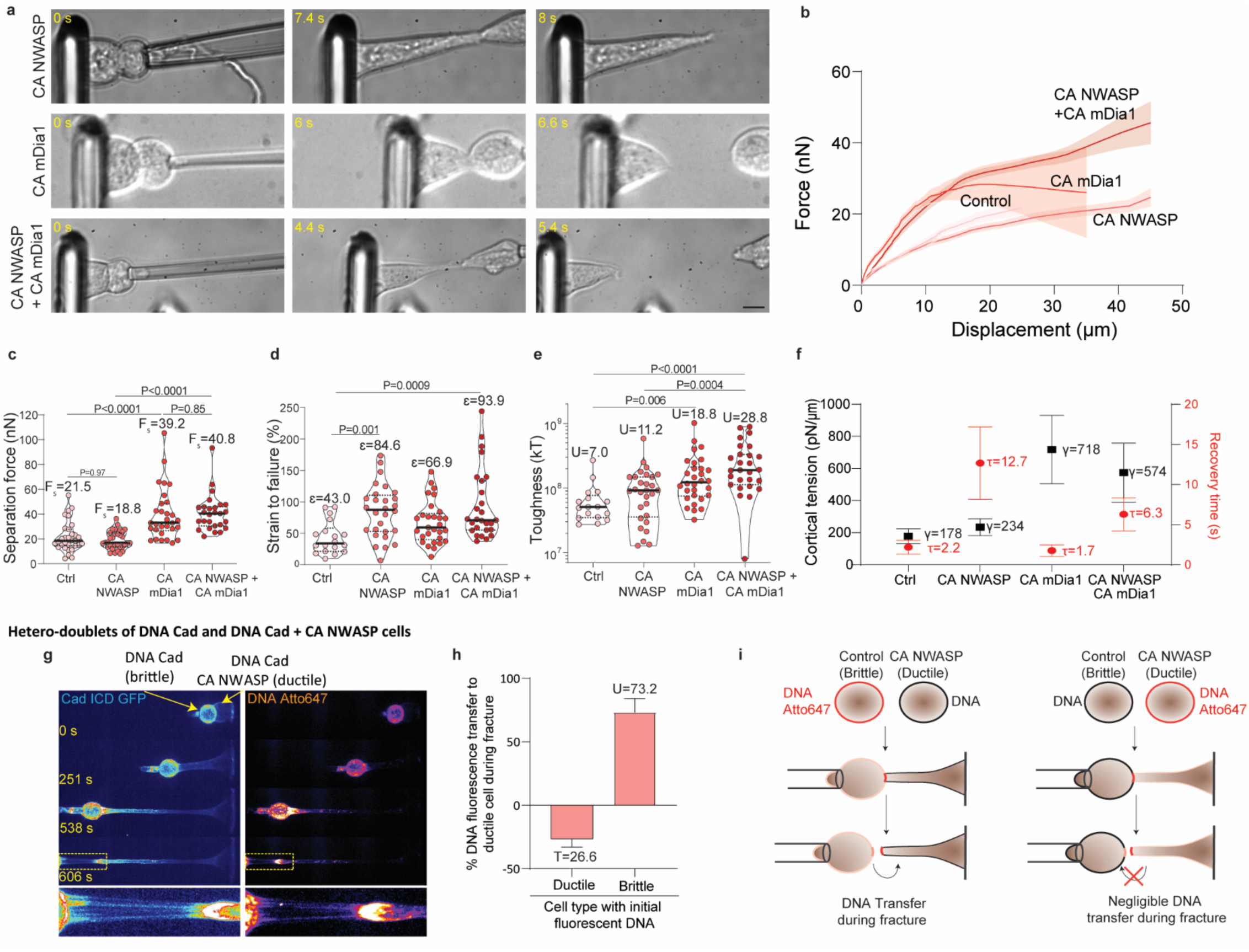
Actin architecture governs viscous dissipation, ductility and hence adhesion toughness. (a) Snapshots of DNA-cad doublets transfected with constitutive active (CA) version of either N-WASP, mDia1 or both during cell separation experiments. Scale 10 μm. (b) Averaged force displacement curves for DNA-cad doublets with either CA N-WASP, CA mDia1 or both (shaded region represents 95% confidence interval of respective curves). (c) Separation force, (d) Strain to failure and (e) Toughness for DNA-cad doublets with either CA N-WASP, CA mDia1 or both, the central line in violin plots represents the median and values indicated above each group represents the mean. (f) Cortical tension (black squares), and characteristic recovery time for cell rounding after fracture (red circles), the point represents the mean and the error bars represent the 95% C.I. (g) Maximum intensity projections of Cad GFP and DNA Alexa fluor 647 signal from a hetero-doublet between DNA-cad (brittle) and DNA-cad – CA N-WASP (ductile) cells pulled to cause junction fracture under nocodazole treatment (10 μM). Scale 20 μm. (h) Relative DNA fluorescence transfer to DNA-cad-CA N-WASP ductile cell after junction fracture under nocodazole treatment (10 μM) in cases when either one of the cell types is labelled using fluorescent DNA (n=8). (i) Schematic depicting separation experiment using differential DNA labelling on hetero-doublets of DNA-cad and DNA-cad + CA NWASP cells lead to transfer of majority fluorescent DNA to the ductile cell irrespective of initial labelling showing that cortex brittleness determines adhesion fracture.

We then asked whether these changes were better correlated with any associated changes in cortical stress or viscous flow. The viscous properties of the cells after fracture were characterised by analysing the shape recovery of the elongated cells after fracture. The characteristic recovery time τ was calculated after an exponential fit. We used τ as a proxy to estimate the viscous property of the cells after they have undergone dissipative deformation during fracture. The results demonstrated more than five-fold increase in recovery time in presence of CA N-WASP (P<0.0001), whereas the cortical tension remains unchanged (P>0.99)(**Figure 4f**). In contrast, mDia1 led to no significant change in recovery time (P>0.89) but almost four-fold increase in cortical tension (P<0.0001). Interestingly the combination of both, led to about three-fold increase in both the recovery time (P<0.0001) and cortical tension (P=0.0002). Based on these results we concluded that an increase in dissipation alone enabled higher ductility; conversely, an increase in tension without a drop in dissipation led to increased force and toughness. Simultaneous increases in both cortical tension and dissipation led to a synergistic increase in adhesion toughness (**Figure 4e and f**). In addition, analysis of the recovery time for increased contractility confirmed that the recovery time corelated well with cell ductility (**Supplementary** Figure 6).

Taken together our experiments strongly suggest that the ductility and brittleness of the cellular contact can be regulated by the balance of actin polymerization and myosin activity. They also suggest that the cohesiveness of cell-cell adhesion results more from the deformability of the cortex than from the binder affinity provided that reciprocal signalling is impeded. The physical mode of separation of brittle junctions (multi-tubular) differed from the elongated conical deformation in ductile cells. However, at the molecular level, it was very difficult to determine how the junction broke. We therefore decided to test heterodoublets of DNA*_45_*-cad and DNA_45_-Cad – CA-N-WASP cells. To identify the localization of the binders after the junction rupture, we coupled one cell type with a fluorescent DNA sequence and the other with the non-fluorescent complementary strand. We subjected the doublet to separation assay in presence of nocodazole. The N-WASP expressing cells deformed much more drastically (hence buffering the effect of nocodazole) than the DNA_45_-cad cells (**Figure 4g**, **Supplementary video 10**) creating a highly skewed deformation of the junction. Upon failure we found most of the junctional DNA fluorescence (73 ± 8%) at the N-WASP cell surface (**Figure 4g-i**) irrespective the initial labelling. **Figure 4g-i** demonstrates that the binders rip off the most brittle cortex and transfer to the ductile one. We concluded that the cortex of the brittle cells is the weakest point of the asymmetric junction. It fractures into numerous tethers, whereas DNA_45_-cad remains bound in trans. The notion that stronger cortices correlate better with higher adhesion strength was further supported by comparing A431D with S180, two different cell lines with distinctly different actin cortex thicknesses (**Supplementary** Figure 7**, Supplementary Text**). Taken together, these data strongly suggest that the deformability of the cortex and the dissipation of energy by its plastic deformation redistributes the mechanical stress at the junction and alters its resistance to mechanical rupture.

### The cortex as an energy sink during adhesion fracture

Cortical actin acts as an energy sink during adhesion failure. Consequently, branched actin networks support better energy storage and dissipation through increased ductility, thus enabling higher adhesion toughness. We therefore postulated that mechanical strengthening of the adherens junction upon E-cadherin recruitment may be associated with regulated changes in cortical properties. We used DNA-cad to test this idea, as this construct abolished the mechanosensitive recruitment of cadherins and their ability to modify the cortex by signalling. We then asked whether these conclusions drawn from the use of DNA-Cad could be extended to bona fide adherens junctions. Given that E-cad can modulate cortical properties, we asked how junctional cohesion changes in E-cad A431D cells upon increased contractility. In contrast to DNA-cad, dissociation force (50%), strain to failure (125%) and toughness (129%) were significantly increased in E-cad cells upon treatment with the highest concentration of nocodazole (10 μM) (**Figure 5a-c**). Surprisingly, we found that E-cad cells treated with 10 μM nocodazole showed more than 2-fold lower cortical tension and f5-fold longer recovery time as compared to DNA-cad cells under the same treatment (**Figure 5d**). The accumulation of stress in actin networks is strongly dependent on their architecture. While mDia1 polymerized networks (linear actin) favour higher force generation and accumulation of myosin generated stress, heavily branched Arp2/3 polymerized networks prevent such stress accumulation ^21^. E-cad can signal through Rac1 and Cdc42 to promote Arp2/3-mediated actin assembly. We therefore asked whether cortical remodelling by this pathway could buffer contractility to prevent an uncontrolled increase in cortical tension leading to a brittle cortex. Indeed, we observed that expression of constitutively active N-WASP in DNA*_45_*-cad cells partially alleviate the ductile to brittle transition observed in DNA-cad cells. Rescue of adhesive properties and ductility for DNA-cad could also be recapitulated by exogenously activating EGFR which is a known activator of Rac1 and Cdc42 and is normally known to be activated by E-cad homophilic ligations (**Supplementary figures 8-10, and Supplementary text**).

**Fig 5.**
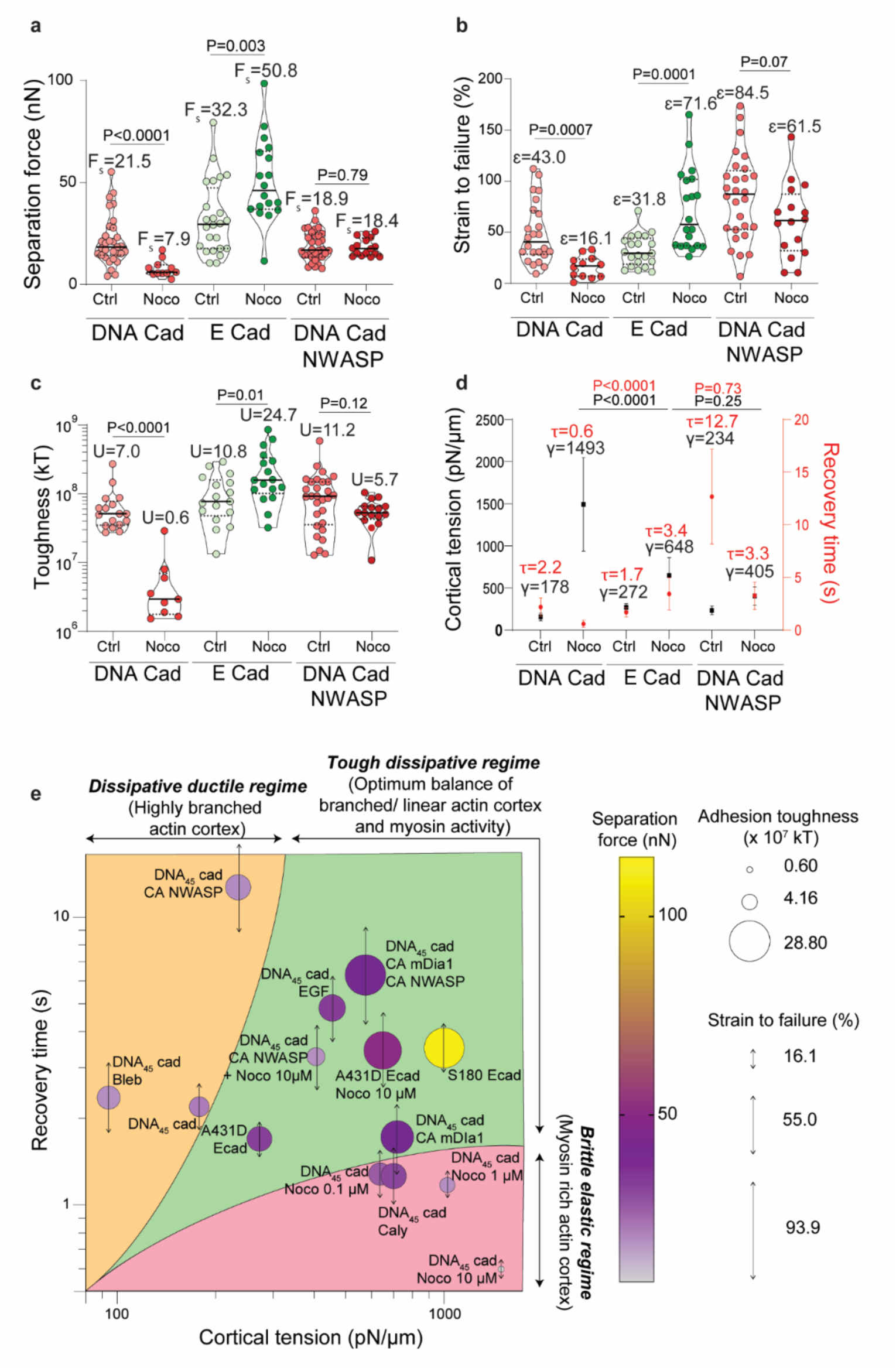
Stiff and dissipative acto-myosin cortex as an efficient energy sink enabling high adhesion toughness. (a) Separation force, (b) Strain to failure, and (c) Toughness for DNA-cad, E-cad, and DNA-cad CA N-WASP doublets under control and nocodazole treatment (10 μM). (d) Cortical tension (black squares), and characteristic recovery time for cell rounding after fracture (red circles), the point represents the mean and the error bars represent the 95% C.I. for DNA-cad, E-cad, and DNA-cad CA N-WASP doublets under control and nocodazole treatment (10 μM). (e) Scatter plot of different cad and treatment types demonstrating presence of different regimes of junction fracture. The x and y axes represent cortical tension and recovery time respectively. Size, color, and double headed arrow length at each point represent adhesion toughness, separation force and strain before failure. The yellow zone closer to y axis is ductile regime where fracture is dominated by the extreme dissipative characteristic of cortex leading to very ductile cells, such a behavior is expected in cortex dominated by branched actin supporting little force generation by myosin. The pink zone closer to x axis is a rigidity dominated regime which leads to increased brittleness, this region is envisioned to be myosin dominated which leads to excessive pre-stress in the cortex and little dissipative character. The central green region is where most wild type systems seem to operate and is a balance of increased contractility and dissipative character, enabling the cortex to efficiently propagate forces and absorb large amount of energy prior to yielding and fracture.

Unlike DNA-cad alone, N-WASP expressing cells showed no significant drop in separation force (P=0.79), strain to failure (P=0.07), and toughness (P=0.12) (**Figure 5a-c**). Further, on treatment with 10 μM Nocodazole, there was only a modest two-fold increase in the cortical tension of CA N-WASP expressing cells, which was not significantly different from the cortical tension of E-cad cells (P=0.25) (**Figure 5d**). In fact, DNA-cad and DNA-cad CA N-WASP cells treated with 10 μm nocodazole showed very distinct behaviour when subjected to micropipette aspiration with the former showing signs of an extremely stiff but fragile cortex with many blebs (**Supplementary videos 12 and 13**).

It demonstrates that the dissipative behaviour (characterized by the shape recovery time) and the rigidity of the cortex (characterized by the cortical tension) collectively govern adhesion toughness. While the former has a deterministic control over ductility, the latter predominantly promotes higher separation force if ductility is maintained. We summarized this by plotting all the experimental observations on a shape recovery time (τ) vs cortical tension (γ) plot (**Figure 5e**). Three possible regimes were identified in the graph: *i*. *Dissipative ductile* characterized by a highly branched cortex which does not support build-up of contractility irrespective of myosin activity and supports high ductility but low separation force; *ii*. *Brittle elastic* with high myosin activity leading to predominantly elastic and brittle cortex; and *iii. Tough dissipative* regime where concomitant increase in tension and recovery time lead to a tough yet dissipative cortex leading to strong cell-cell adhesion.

### Mechanical modelling

To validate the interpretation of our results, we have constructed a descriptive mechanical elasto-plastic model for the separation of a cell doublet. For simplicity, we take advantage of symmetry and model the cell doublet as a single cell, considered as a fluid-filled capsule adhering to a flat substrate. We performed numerical calculation of cell detachment using the model described in **Supplementary Text**. Briefly, the cell cortex is considered locally elasto-plastic with a non-linear constitutive stress-strain relationship. Myosin activity generates an isotropic active stress that can be tuned to increase the cortical tension of the capsule. It adds to the anisotropic stress components generated by the external strain exerted by the pipette on the cell pole opposite the junction. We also considered the viscous flow generated by the gradient of elastic stresses advecting the binders and myosin. We introduced an adhesion energy per filament and a threshold for the mechanical energy above which the filaments break. A full description of the model can be found in **Supplementary text**.

We first tested whether our model successfully describes the initial ring-like distribution of the cadherin and actomyosin complex. The junction architecture at equilibrium showed a homogeneous distribution of both components by a simple minimisation of the energy for a contact initiated between two homogeneous cortices. However, when we introduced a local depletion of the cortex at the centre of the contact, both the binders and the acto-myosin cortex relocalised to the edge of the contact (**Figure 6** **a,b**). Consequently, the mechanical stress distribution generated by myosin also peaked at the edge of the contact (Figure 6c). Increasing cortical tension resulted in a monotonic increase in junction diameter and an increased localisation of binders at the rim. Thus, the simulated results mirrored the experiments.

**Fig 6.**
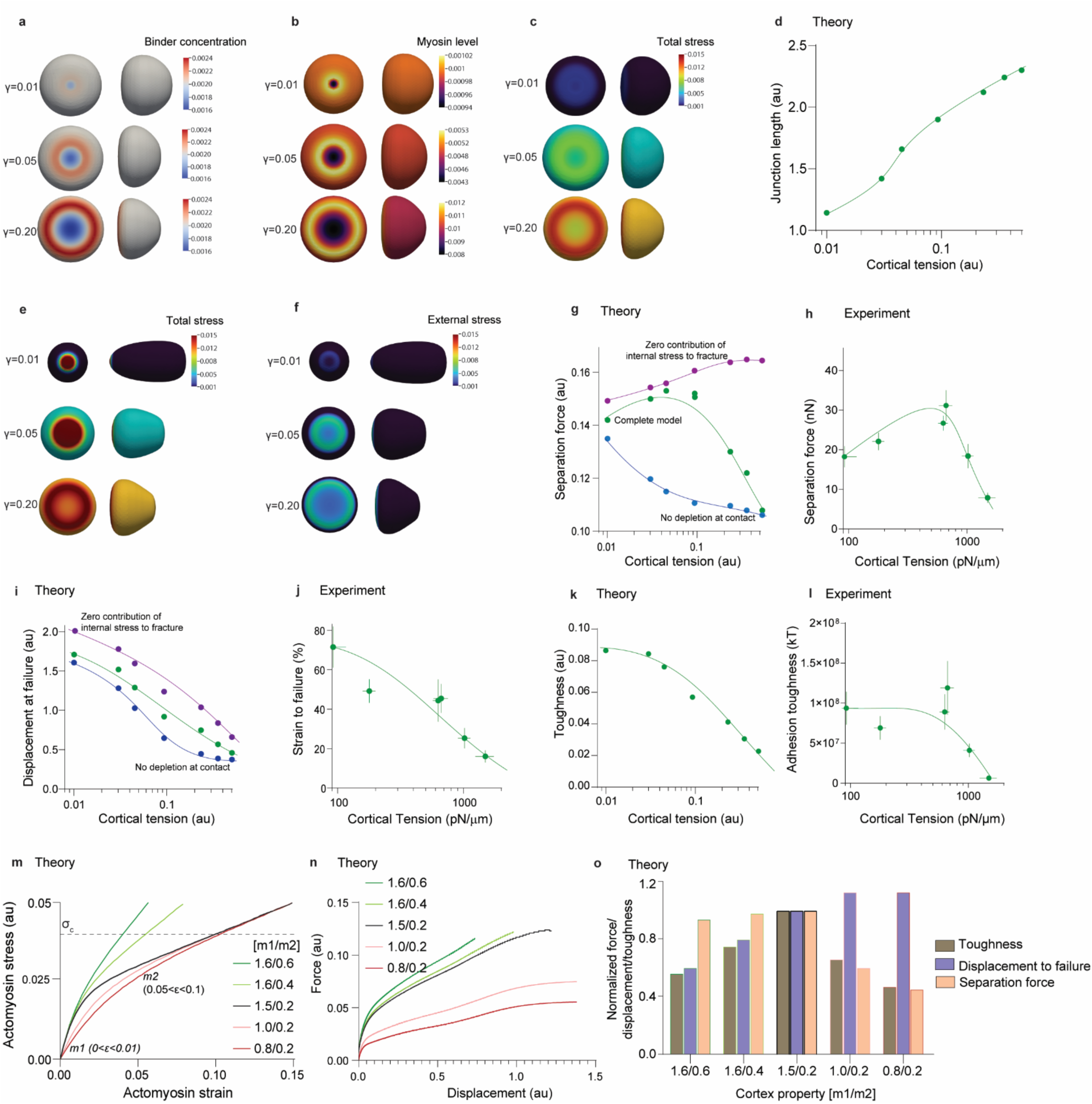
Computational model for cell adhesion fracture predicts that elasto-plastic properties of the actin cortex are deterministic of cell adhesion strength. Organization of adhesion in terms of (a) binder, (b) myosin, and (c) total stress distribution immediately after adhesion formation and prior to application of external force at different contractility. (d) The model recapitulates contractility dependent expansion of cell-cell contact as junction length correlates well with cortical tension. (e) and (f) Spatial distribution of total stress and externally applied stress immediately prior to adhesion fracture in cells with different contractility. (g) Separation force as predicted by the complete model (green), model ignoring contribution of myosin generated force towards cortex rupture (violet), and model where initial outward flow in the contact plane of binder and cortex is not incorporated (blue). (h) experimental values of separation force for DNA-cad. (i) Displacement to failure as predicted by the complete model (green), model ignoring contribution of myosin generated force towards cortex rupture (violet), and model where initial outward flow in the contact plane of binder and cortex is not incorporated (blue). (j) experimental values of strain to failure for DNA-cad. (k) and (l) toughness values as a function of cortical tension as predicted by the model and observed in experiments respectively. (m) Stress-strain curves for five different cases used to simulate cell cortices with different inherent elasto-plastic characteristics, *m1* and *m2* are slopes of the curves for each group at low (0<ε0.01) and high (0.05<ε0.1) strains representing the elastic and plastic regions respectively. The dotted line represents the critical stress (σ_c_) at which the cortex fractures. (n) Force displacement curves for the five different cases in (m) when subjected to pulling till fracture of adhesion. (o) Comparison of separation force, displacement to failure and toughness for the five cases with different elasto-plastic character confirms that a cortex with high stiffness and plastic character enables best performance during adhesion fracture.

In a first set of simulations, we tested the effect of myosin activity on junction separation while keeping the strain-tension relationship of the cortex constant. We simulated the pulling experiments (**Supplementary Video 13**) under different levels of internal stress generated by different levels of myosin activity (increased cortical tension). **Figure 6e-f** shows the distribution of the total mechanical stress and its external component just before reaching the rupture point for different cortical tensions. Increasing the cortical tension decreased the global deformation of the cell at rupture, whereas decreasing the cortical tension increased the deformation of the cell and caused the junction to shrink before rupture. We calculated the separation forces Fs in different cases. We considered applying the threshold for the bound rupture using either the externally applied stress or the total stress (external and myosin generated). We also compared junctions with or without central depletion of cortical actin. **Figure 6g-h** shows that we were able to recapitulate the observed non-monotonic variation of separation force with the increase in cortical tension only when we considered the total stress at junctions presenting cortex accumulation at their edge. According to the model, at low contractility, the externally applied stress dominates the internal one. The external force when born by smaller junctions generates a higher local stress. In this regime, the increase in dissociation force with cortical tension is mainly due to the increase in junction size. At high cortical tension, the internal stress becomes more dominant, with the external stress providing only an excess of mechanical tension necessary to reach the fracture point. As the stress generated by myosin scales with junction size and as junction size plateaus, the amount of external force required to break the junction decreases as the internal stress increases (brittle regime under high cortical tension). Oppositely, the stress accumulated during the junction rupture accumulated quite uniformly along the free cortex independently of the assumptions we considered It resulted that ε decreased monotonically with the increase of cortical tension (**Figure 6 i-j**) line with the observations. Consequently, the toughness U at low cortical tension plateaued for the lowest cortical tension to subsequently it decrease at high cortical tensions. This trend properly accounts for the experimentally observed behaviour (**Figure 6 k-l**). We concluded that our model adequately described the ductile to brittle transition following increase in cortical tension in absence of actin regulation.

In the second set of simulations, we held myosin activity constant and varied the constitutive strain-stress relationship of the actin cortex. We assumed a plastic behaviour of the cortex (**Supplementary Text**) with a viscoelastic coefficient m_1_ before a critical yielding deformation and a viscoelastic coefficient m_2_ after. This is a crude approach to account for changes in the amount of bundled and branched actin. As our experiments are performed at a constant strain rate, the values of each parameter m_1_ and m_2_ comprise an elastic and a viscous contribution. The plastic deformation accounts for a change in cortex behaviour at low *vs* high strain as reviewed in ^22^. We used a fixed local criterion for bound rupture that we apply to the total stress in all simulations. **Figure 6** **m** shows the different constitutive relations we used. We first Increased m_1_ (from 0.8 to 1.5) and kept m_2_ constant (0.2). It reflects an increase of the viscoelasticity at small deformations only (linked to an increase in elasticity and/or viscosity) (**Figure 6 n-o**). The separation force increased as well as the toughness whereas the displacement slightly decreased. Then, We held m1 constant(1.5) and we increased m_2_ (from 0.2 to 0.6). The toughness and the displacement to failure consequently decreased whereas the force to rupture hardly changed. The calculation showed that enhancing the cell plasticity (smaller m_1_) enabled the propagation of strain further along the cortex precluding build-up of large stress at the junction and dissipating energy along the cell cortex (**Supplementary video 14**). It demonstrates that the contact strength, defined as the set of all three values for the Force at rupture, maximal strain and toughness, is a compounded property depending on the non-linear rheological behaviour of the whole cells and cannot be inferred from the local property of the junction itself. The total deformability of the cell must be accounted for since it contributes to localising stress at the junctions. It strongly reinforce the idea that a the simple value of the cortical tension and density of binders is not sufficient to describe the junction strength. Phase diagrams like the one we established in **Figure 5** are required to better describe how a contact resist to mechanical load. We propose that the relaxation time τ of the cell shape is a practical (though not completely physically accurate) way to account for the viscoplasticity of the cell cortex. It is consequently a handy proxy to understand the changes in the mechanical behaviour of the contact. How the molecular balance (and its regulation) between branched and linear actin quantitatively controls the deformability of the cells remain to be uncovered.

## Discussion

While previous studies have mainly explained cell adhesion in terms of cortical tension and binder affinity ^6, 8^, here we show that cortical ductility, which is in turn determined by cortical architecture, plays a central role in determining cell adhesion strength. Early studies explained the preferential homophilic adhesion between cadherins based on differential affinities between these molecules at the single molecule level; however, the magnitude of the difference in molecular binding strength does not explain the large differences in cell adhesion strength between E-E, N-N and E-N cad expressing cells ^9^. Our data confirmed that in the absence of signalling from cadherin molecules, even very large differences in the affinity of the binding molecules do not mediate a change in cell adhesion strength (**Figure 2**). In contrast, modulation of the cortex by signalling such as EGFR strongly influences cell adhesion strength. EGFR signalling, which can be activated by homophilic ligation of cadherins, is known to activate Rho GTPases, namely RhoA, Cdc42 and Rac1, thereby altering the architecture and biophysical properties of the actomyosin cortex. ^14, 16, 23, 24^. These Rho GTPases are differentially activated by homophilic ligation of different cadherins, which may explain the different strengths they exhibit.^25^. Interestingly, as previously reported for cadherin expression levels, we saw a strong influence of DNA-cad number on cell adhesion strength. Unlike cadherins, DNA-cad ligation did not modulate cortex properties via signalling, so the origin of this influence is unlikely to be a result of receptor number-dependent signalling. However, this points to the importance of stress redistribution in the cortex, as a higher density of tethers may promote better stress redistribution in the cortex, allowing it to withstand higher overall forces.

The formation of tethers during the rupture of cell-cell adhesions has been noted previously ^8^ as well as in the present work (**Figure 3**). Since the breaking of cadherin trans-interactions is the last step, this clearly indicates that deadhesion proceeds from the inside out. A simple model would be to consider the adhesion machinery as a series of springs (cad-cad, cad - β-cat, β-cat - α-cat, α-cat - F-actin bonds) with finite breaking stresses and to identify one of them as the weakest link. However, this is extremely unlikely given the diversity of adhesive strengths that can be achieved using exactly the same links (**Figure 5**). Such a scenario also does not explain the consistent failure of the intracellular linker apparatus in control DNA-cad cells compared to DNA-cad-CA-N-WASP cells in hetero doublets, as all the above linkers remain the same in both cases. However, they can be explained if the origin of the rupture lies in the cortex, especially considering the diversity that exists in the cortical architecture in terms of thickness, branching and connectivity. ^26^. Therefore, we propose that the fracture is initiated in the cortex itself and its propagation to the junction depends on the cortical architecture. For example, a highly pre-stressed cortex (**Figure 3**, 10 μM nocodazole) will support little deformation and cracks will propagate rapidly, leading to catastrophic failure at low external stresses and strains, as evidenced by the formation of numerous tethers. On the other hand, a cortex with a mixture of stressed and deformable elements will allow stress transfer to residual deformable elements as micro-fractures initiate in pre-stressed elements, thus allowing greater deformation while maintaining high forces (**Figure 4**, CA NWASP + CA mDia1). A similar mechanism has previously been described for double network hydrogel materials, where the combination of a highly cross-linked brittle network with a loose ductile network acts synergistically to prevent macroscopic fracture propagation and produce extremely high toughness. ^12, 13^. The presence of two distinct but intertwined actin networks in the medio-apical cortex of Drosophila epithelium undergoing apical constriction is consistent with this notion, where the persistent network serves to maintain epithelial integrity and force propagation, while the pulsatile network acts as a sacrificial actin network during the contractile phase ^27^. Furthermore, it has been shown that F-actin turnover on a timescale consistent with the contractile impulse is essential to maintain force balance and allow rapid reattachment of the actomyosin network to the AJ. This turnover is critical for disassembling the contracted (sacrificial) medial F-actin and rapidly forming a more homogeneous apical network^28^. It can therefore be concluded that the strength of cell-cell adhesion is largely regulated by modulating the structure of the actomyosin cortex, rather than by the strength of the interaction between any two proteins in the cortex.

The cortex can therefore be thought of as an energy sink during the process of cell-cell adhesion. Energy storage or dissipation can occur in a number of ways, namely: (i) stored elastic energy, which is released during rapid rounding of cells after fracture ^9^; (ii) Energy dissipated by the realignment of actin filament ^29^; (iii) Energy dissipated as heat when some actin filaments are severed under the external load; and (iv) Energy dissipated when active turnover of F-actin takes place consuming biochemical energy^28^. While the last factor in the above list is expected to contribute minimally during rapid deadhesion (<5-10s) such as in this work, its role may become exceedingly large at longer time scales as shown previously through stress relaxation experiments on epithelial cell sheets^30^.

It is noteworthy that E-cad modulates the cortex in such a way as to increase cortical tension while maintaining its deformability or ductility. It is important to note that cortical tension is an aggregate property that is a function of myosin activity, F-actin content and architecture. While AJ formation led to an increase in cortical tension, its presence also buffered an uncontrolled increase in stiffness and brittleness, unlike DNA-cad mediated junctions (**Figure 5**). The ability of CA NWASP to partially rescue nocodazole-induced brittleness and buffer the uncontrolled rise in cortical tension supports a strong role for the balance between different Rho GTPases in controlling cortical properties. Furthermore, as signalling strongly controls different actomyosin modulatory pathways, it allows cells to precisely control adhesion strength over a very wide range, thus providing inside-out control of cell-cell adhesion. Overall, it can be concluded that the architecture of the actin cortex controls the viscous dissipation, ductility and hence the strength of cell-cell adhesion.

## Acknowledgements

Work in Australia was funded by grants and fellowships to ASY from the NHMRC (GNT1136592, 1163462) and ARC (DP220103951) and to IN from the European Molecular Biology Organization (EMBO ALTF 251-2018). VV acknowledges the support from the NRF (NRF-NRFI2018-07-00) and from the Mechanobiology Institute seed funding

## Materials and Methods

### Plasmids and reagents

All cadherin-based constructs were derived from full length mouse E-cad cloned in pEGFPN1 vector between *Hind*III and *Age*1 sites. For Halo modified cadherin, extracellular domain was deleted and Halo tag was inserted between signal peptide and transmembrane domain of E-cadherin. Further, to reduce endocytic uptake of modified DNA-cad receptors, three point mutations in the cytosolic portion of cad using site directed mutagenesis were carried out (K740R, L743V, and L744V)^19^. Constitutively active version of N-WASP was generated by performing four point mutations (K183E, K186E, K189E, and K192E) as reported previously^31^. The constitutively active version was then fused at the C terminal of Halo-E-Cad-GFP sequence. For constitutively active version of mDia1, C terminal fragment of mDia1 (753-1255), i.e., without the Diaphanous inhibitory domain was cloned at the c terminal of blue fluorescent protein in pIRES-Hyg2 vector.

Oligonucleotides with 5’ HaloTag Ligand(O4) modification with or without 3’ Atto647N(c) modifications were procured from biomers.net GmbH. The sequence for 45F, 45R and 12R oligos were 5’-GGCTCCCTTCTACCACTGACATCGCAACGGATCTTAACGCACTTG-3’, 5’-CAAGTGCGTTAAGATCCGTTGCGATGTCAGTGGTAGAAGGGAGCC -3’, and 5’-CAAGTGCGTTAA-3’ respectively.

### Cell culture, transfection and cell line generation

Routine cell culture was performed in high-glucose DMEM, supplemented with 10% FBS. For all live imaging experiments, phenol red free DMEM was employed. All transfections were done using Lipofectamine 3000 reagent as per manufacturers protocol in 24 well plates. A ratio of 1:2 for plasmid DNA to Lipofectamine/Power Reagent was used with 1 μg plasmid DNA per well of a 24 well plate. For making stable lines, cells were selected and maintained in Geneticin (500 μg/mL) or hygromycin B (200 μg/mL) after transfection. For experiments, cells were used within 2-4 population doublings after FACS sorting to enable a narrow range of expression of protein of interest.

### Cell aggregation

For cell aggregation of E-cadherin expressing cells, the cells were dissociated using enzyme free cell dissociation buffer and re-suspended in DMEM supplemented with 1% Bovine Serum Albumin (BSA). This suspension was incubated at 37°C for 45 mins for the formation of cell doublets and bigger aggregates.

For DNA-cad expressing cells, the Halo tag expressed on the cells was first allowed to react with Halo ligand modified ssDNA by addition of oligonucleotides to cell culture medium at final concentration of 1 nmole/ml. Cells were then washed with 1X PBS and dissociated using enzyme free cell dissociation buffer. Equal number of cells labelled with complementary DNA strands were then mixed and resuspended in DMEM supplemented with 1% BSA and 25 μM para-amino-Blebbistatin. The cells were incubated at 37°C for 45 mins to allow for cell-cell adhesion after which the cells were centrifuged, washed and re-suspended in fresh medium.

### Immunoblotting

Cells post aggregation and appropriate control (EGTA for E-cad and Cells without DNA labelling for DNA-cad) were incubated on ice with RIPA lysis buffer (Sigma Aldrich) supplemented with proteases and phosphatases inhibitor coc_kbT_ails (Sigma Aldrich). Lysates were then subjected to SDS-PAGE, transferred to nitrocellulose membranes, and incubated with relevant antibodies. The antibodies used were pEGFRY845 (44784G, Thermo Fisher Scientific), EGFR (2223, Cell Signaling Technologies), pMLC2 (3674, Cell Signaling Technologies) and β-actin (MA515739, Thermo Fisher Scientific). Immune complexes were detected with appropriate HRP-conjugated secondary antibodies from Cell Signaling Technologies and enhanced chemiluminescence reagent (Clarity ECL, BioRad). Protein band intensities were quantified using ImageJ Software.

The human RTK phosphorylation assay was performed using an antibody array membrane accordingly to the manufacturer specifications (ab193662, Abcam).

Spots intensities on the arrays were quantified with ImageJ Software.

### Cortical tension measurement

A medium filled chamber with side access for glass micropipette was devised using two pluronic F127 treated (1%; 1 hour) glass coverslips (24 mm x24 mm, 24 mm x 55 mm), and a custom made U shaped PDMS gasket (3 mm high). The chamber was filled with a cell suspension and cells were subjected to micropipette aspiration, using passivated glass micropipettes (5-7 μm diameter, Biomedical-Instruments, Germany). Cortical tension was calculated using the law of Laplace:

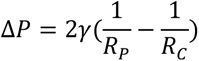

Here, R_P_ is the radius of the pipette, R_C_ is the radius of the cell and ΔP is the pressure when the aspirated portion of the cell was a hemisphere of radius R_P_. The zero pressure in pipette was calibrated prior to each experiment run by validating lack of inward or outward movement of floating debris/particles in the media at the tip. Thereafter, pressure is decreased gradually till the aspirated cell forms a hemisphere at the pipette tip.

### Cantilever microfabrication

The cantilevers are fabricated on silicon wafers as substrate for a two-layers aligned lithography, then transferred to their final substrate, a glass coverslip, using an optical UV-curable adhesive as the transfer medium.

Here we describe the detailed procedure, Extended Fig. 3a shows a schematic of the principal steps. A standard Si wafer, 4” diameter and (100) orientation is de-hydrated on a Hot Plate at 200 °C for 10 min. After letting it cool to room temperature, a layer of PMGI SF6 (Microchem Newton, MA 02464, USA) is spin coated at 1500 rpm for 45” and baked at 190 °C for 1 min, giving a layer of about 600 nm thickness (Ex. Fig. 3a, i). On top of the PMGI layer, we then spin-coat the first layer of SU-8 (Microchem): we use SU-8 3050 at 2700 rpm which after soft-baking for 20 min at 95 °C results in about 50 µm thickness. This SU-8 layer is exposed for 10” to UV light through a photomask with the lay-out of the cantilevers (Ex. Fig. 3b and layer 1 of the provided gds file with the designs of the two masks). For the exposure we use I-line filters on a MJB4 mask aligner (SUSS MicroTec, 85748 Garching, Germany), the light intensity measured after the filters is 23 mW/cm2. Post-bake of the exposed resist is then applied on a Hot Plate at 65 °C for 1 min followed by 3 min at 95 °C (Ex. Fig. 3a, iii). On the post-baked resist, we then spin-coat a second layer of SU-8, SU-8 3050 at 2100 rpm for 45”. The wafer is then placed on a Hot Plate at 65 °C and then the temperature is slowly raised up to 95 °C. The temperature ramping-up takes about 20 min, then after keeping the wafer 20 min more at 95 °C the Hot Plate is switched off and allowed to cool down to room temperature naturally. The second layer is then exposed with the optical mask with the second lay-out using the same parameters as for the first exposure (Ex. Fig. 3a, iv). Post-baking on the Hot Plate for 1 min at 65 °C followed by 3 min at 95 °C is done, with slow ramping-up of the temperature (same as before) and slow cooling down at the end (natural cooling, same as before). At this stage, the resist is developed by immersion in SU-8 Developer (Microchem) for about 10 min. The final total thickness of the features is 120 µm (50 + 70 µm) (Ex. Fig. 3a, vi). A glass coverslip is now prepared as the final substrate for the cantilevers. We use 1.5 square coverslips, 25 mm in length. Before usage, they are cleaned first in an ultrasound bath of soapy water for 5 min, then after a first rinsing in clean DI water they are washed with Acetone and IPA and finally dried with a N2 blow gun. On a clean coverslip a layer of UV-curable optical glue (NOA 73, Norland Products Inc., Jamesburg, NJ 08831, USA) is spin coated and pre-cured with a short exposure to UV light. We use an UV-LED system (UV-KUB2, Kloé 34270 St Mathieu de Tréviers, France) with 354 nm UV light and 25 mW/cm2 power density; exposing for 30 s the spin-coated layer (2000 rpm for 45 s, final thickness about 30 µm) the NOA73 becomes solid enough to avoid reflow but it retains enough adhesion power for the following step (Fig 1a, vii). The pre-cured NOA73 is pressed in contact with the developed SU-8 features, then a second exposure to UV light in the UV-KUB system (1 min at 25 mW/cm2 power density) complete the curing of the glue and promote the adhesion of the SU-8 to the coverslip. Finally, the cantilevers are released by dissolving the PMGI in a water solution of potassium borate, commercially available as photo-resist developer AZ 400K (AZ Electronic Materials, Somerville, 08876 NJ, USA). Dissolving the PMGI releases the SU-8 cantilevers from the silicon substrate and leaves them attached to the coverslip thanks to the cured NOA73 glue (Fig 3a, viii). After successful detachment of the Si substrate, the developer solution is removed and quickly replaced with DI water. The suspended cantilevers, attached to the coverslip, are kept wet in clean DI water to avoid collapsing, until they will be used. Such device can be stored safely for months.

### Cell separation assay

For cell separation assays, the cantilever microchips were coated with poly-L-Lysine (0.1%), overnight at 4 °C, followed by bovine collagen type I (0.2% in 0.2% acetic acid) (Corning life sciences, USA). The cantilevers were washed with 1X PBS and transferred into the custom setup made of 60 mm petridish with a 2 cm section of vertical wall removed, a glass coverslip and U shaped PDMS gasket (5 mm height) in between the two as depicted in Supplementary Figure 11a. The reservoir thus obtained was filled with DMEM supplemented with 1% BSA. Cell suspension containing cell doublets was then gently added to the front reservoir region of the chamber. Under the microscope, cell doublets were then captured using the micropipette connected to micromanipulators (Narshige, Japan) and were then positioned at the edge of one of the cantilevers. The cell away from the pipette tip was then allowed to adhere to the cantilever for 8-10 mins. Finally, the other cell held by the micropipette was pulled away using the micromanipulator until the two cells were separated. At any given time, force applied was quantified by measuring the deflection of cantilever tip (Fig. 2a). In addition, strain on cells and displacement of pipette tip relative to cantilever was alsuppl

so measured. For measurement of recovery time, the cell aspect ratio was measured starting from the point of fracture and exponential was fit with plateau value being set at 1. The characteristic time τ was then calculated from the curve using GraphPad Prism software.

### Cell separation with confocal imaging

For confocal imaging during cell separation, a modified setup as depicted in Supplementary Figure 11b was employed. Herein a 1 mm high collagen coated PDMS strip was bonded to a glass coverslip which was then treated with Pluronic F127 (1%, 1 hour) to prevent cell attachment. The reservoir was then filled with cell suspension. Like the cell separation assay cell doublets were captured using pipette and the free cell was allowed to adhere to the PDMS strip, increasingly large displacements were then applied in a step wise manner using micromanipulators and a z stack was acquired using spinning disk fluorescence microscope with Metamorph software at each of the pulling steps.

### Statistical analysis

All statistical analysis, and graphs were plotted using GraphPad Prism software. For statistical analysis data were first tested for normal or lognormal distribution in using KS test. In case of normal distribution, Students t-test was performed for comparison between two groups and ANOVA with Tukey’s post hoc tests were done for more than two groups. The same was done after log transformation of data in case of lognormal distribution. Non-parametric test Mann-Whitney test was performed where data failed KS test for both normal and lognormal distribution. Individual details on tests performed, P values, number of independent experiments and data points for each experiment have been included in the supplementary text.

Detailed methods for the theoretical model are included in the supplementary text.

## Notes

### Competing Interest Statement

The authors have declared no competing interest.

