## Supplementary info for "Cortical ductility governs cell-cell adhesion mechanics"

Affiliations:

### Model formulation

#### Basic assumptions

We consider the cell to be a fluid filled axisymmetric capsule [1] where the surface of the capsule wall represents the cell membrane and cytoskeleton. In order to keep the model simple, the cell is assumed to adhere to a flat rigid substrate instead of another cell which in the case in the experiments. The model presented here can, however, be extended to inter-cellular adhesion as well. Elastic forces within the cell are due to the bending and in-plane stretching stiffness of the cytoskeleton and cell membrane combined [2]. The DNA linker driven cell-substrate adhesion forces are also taken to be elastic in nature (more details below). Since the cell detachment due to the forces by the pipette takes place at a very slow rate (?? mm/min) we consider the detachment process to be quasistatic. The quasistatic assumption further results in another simplification that we do not have to consider any hydrodynamic interactions within cytoskeleton and other parts of the cell and external fluid. We also consider the resistance of the cell against change in volume but since is cell surface is not considered impermeable the cell volume is not observed to be conserved in the experiment [REF]. We assume that the adhesion of the cell to the substrate takes place over a long period of time. During this process the cytoskeletal remodeling takes place and the cell reaches a new mechanical equilibrium. The cytoskeletal remodeling during the adhesion process is assumed to be primary by the flow of cytoskeleton along with the myosin proteins and DNA linkers along the cell surface [3, 4]. On the other hand the cell detachment is considered to take place at a relatively shorter duration (but still at a slow rate to ensure quasistatic assumption). Therefore, during the detachment the cytoskeletal remodeling is not considered. Further, we also do not assume any source of production and degradation of actin, myosin and DNA linkers making their respective quantities fixed in the adhering cell [5].

#### Geometry

The cell surface is considered as a smooth and continuous axisymmetric surface as shown in figure below. The axisymmetric description reduces the cell geometry to a two-dimensional curve parametrized by $s_0$ in the initial configuration [6]. Here we denote all the geometric quantities in the initial configuration with a subscript $_0$ and in the adhered or partially detached state they are denoted without the subscript.

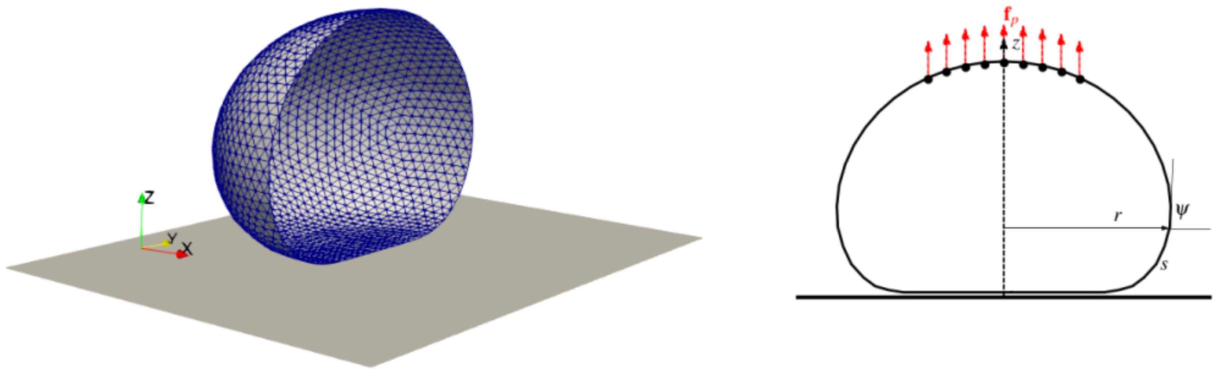


Figure 1: Schematic showing cross-section of a cell in adhered configuration to a rigid plane. The red arrows in the right panel denote the forces applied by the pipette.

In the adhered configuration the surface area and volume of the cell are, respectively, given by

$$A_0 = \int 2\pi r_0 ds_0$$
$$V_0 = \int \pi r_0^2 \sin\psi_0 ds_0 \tag{1}$$

where $r_0$ is the radial location of the cell surface and $\psi$ denotes the tangent angle.

Since we are considering the cytoskeletal flow along the cell surface we need to describe the viscous forces due to this flow which is given in terms of the surface gradient (see details below). The surface gradient operator is given by [7]

$$\nabla_s = (\mathbf{I} - \mathbf{n}\otimes\mathbf{n})\,\nabla \tag{2}$$

where $\mathbf{I}$ is the identity tensor and $\mathbf{n}$ is the outward unit normal on the cell surface.

Application of the external force due to cell adhesion and due to the pipette pull results in changes in the shape of the cell along with stretching of the cell surface. This deformation also results in change of the arc length parameter from $s_0$ in the adhered configuration to $s$ in partially adhered configuration. We describe the distribution of the actin network, myosin and DNA linkers along the cell surface in terms of their respective densities. In general these densities can be described in terms of either $s_0$ or $s$ but here we take *material point* $s_0$ as the independent geometric parameter to describe the densities. The density of the actin network along the cell surface is described as a scalar density $\rho_c(s_0)$ in the adhered configuration. Similarly the densities of myosin $\rho_m(s_0)$ and adhesion DNA linkers $\rho_a(s_0)$ on the cell surface are also described. Since there is no production and degradation of actin, myosin and DNA linkers on the cell surface we have

$$\frac{1}{A_0}\int \rho_c dA = \rho_c^0 \tag{3}$$
$$\frac{1}{A_0}\int \rho_m dA = \rho_m^0 \tag{4}$$
$$\frac{1}{A_0}\int \rho_a dA = \rho_a^0 \tag{5}$$
$$\tag{6}$$

where $A_0$ is the cell surface area before adhesion, $\rho_i^0$ for $i \in \{c, m, a\}$ represent the global concentrations of actin, myosin and DNA linkers in the cell and integration is performed on the cell surface. These three quantities are taken as independent parameters in the numerical simulation of the model.

#### Mechanics

In this description there are three internal sources of mechanical response of the cell- passive mechanical response of the actin network [8], resistance against volumetric and surface area changes in the cell, and active forces due to myosin [9]. The mechanical response of the cell is against two external forces - force due to pipette and adhesion forces between cell and substrate. In the following we will describe these constitutive model ingredients in detail.

##### Passive mechanics of actin network

In order to describe the constitutive laws for the passive response of the actin network we consider it to be a homogenized material where the microscopic length scale of the actin network is much smaller than the length scale of the cell [REF]. In this continuum framework for actin network we consider its passive response to be nonlinear viscoplastic at low to moderate stress values [REF]. In order to incorporate the fracture of the cytoskeleton during cell detachment we also assume that actin network fractures when the mechanical stress crosses a pre-specified threshold value (to be described in details below).

In the passive viscoplastic response of the actin network we consider the total stress to be composed of elastic $\boldsymbol{\sigma}_e$ and plastic or viscous $\boldsymbol{\sigma}_v$ components as [REF]

$$\boldsymbol{\sigma}_p = \boldsymbol{\sigma}_e + \boldsymbol{\sigma}_v. \tag{7}$$

In this description with plastic flows the stress-strain response of the actin network is expected to demonstrate a strain softening behavior [10]. This strain softening behavior can be attributed to the plastic flows in the cytoskeleton. We model the elastic contribution of the actin cytoskeleton response by considering it a hyperelastic material with

$$\boldsymbol{\sigma}_e = k_e \left(\alpha_1 \boldsymbol{\epsilon}_e + \alpha_2 - \alpha_2 \exp\left(-\boldsymbol{\epsilon}_e/\epsilon_y\right)\right) \tag{8}$$

$k_e \left(\alpha_1 + \alpha_2/\epsilon_y\right)$ is the elastic stiffness of the actin network for very small strains, $\boldsymbol{\epsilon}$ is the cytoskeleton strain, $\alpha_i$'s are material parameters, and $\epsilon_y$ is the strain corresponding to the onset of yielding. This choice of the constitutive relation ensures a strain softening elastic response for $\alpha_i > 0$ for $i \in \{1, 2\}$. We will discuss the specific choice of values for $\alpha_i$'s in a separate section below.

For the viscous response of the actin cytoskeleton we model it a gel with constant viscosity [11, 12]. By assuming very slow flow rates on the cytoskeleton along the cell surface ($R_e \approx 0$) we can describe the cytoskeletal flow as Stokes flow and write

$$\boldsymbol{\sigma}_v = 2\mu \nabla_s \mathbf{u} \tag{9}$$

where $\mu$ is the cytoskeleton viscosity and $\mathbf{u}$ is the velocity field. It needs to be noted here that thanks to the polarity of F-actin the actin cytoseleton is intrinsically a polar medium [11] here we are ignoring that polarity from the cytoskeletal description and considering a homogenized state of the network where F-actin of different polarities are isotropically distributed .

#### Active force due to myosin

As mentioned earlier the distribution of myosin in the cytoskeleton is described in terms of its density $\rho_m$. The active forces due to myosin are taken into account in the form of active isotropic contractile stress [13]

$$\boldsymbol{\sigma}_m = \zeta \frac{\tilde{\rho}_m}{1 + \tilde{\rho}_m} \mathbf{I} \tag{10}$$

where normalized myosin density $\tilde{\rho}_m = \rho_m/\rho_m^0$ takes into account the saturation in the active forces at very high myosin density. Here $\zeta$ is the effective coefficient which depends on the cytoskeleton viscosity, actin filament length, myosin velocity along actin filament and unbinding time of the myosin [11].

It has to be noted that similar to the passive actin response here also we have ignored the polarity of the F-actin in the network under the assumption of isotropic distribution of the cytoskeletal filament. With these descriptions of the passive and active contributions of the cytoskeletal response the total stress in the cytoskeleton can be written as

$$\boldsymbol{\sigma} = \boldsymbol{\sigma}_e + \boldsymbol{\sigma}_v - \boldsymbol{\sigma}_m. \tag{11}$$

This internal mechanical stress in the cytoskeleton is balanced by the external forces arising from cell-substrate adhesion, and pipette displacement.

#### Cytoskeletal fracture

In order to take into account the experimental observation of cytoskeletal fracture during cell detachment we consider cytoskeleton to undergo fracture when maximum in-plane stress crosses a pre-specified threshold $\sigma_c$. To estimate the maximum stress at any point on the cell surface we calculate the largest principal stress (magnitude and direction). This stress value at all the locations is then compared with $\sigma_c$ to identify the locations of cytoskeletal fracture. The fracture of the cytoskeleton at any location on cell surface results in loss of the DNA binders at that location.

#### Adhesion forces

As described in the experimental setup the adhesion of the cell is via DNA linkers. In the model we consider the DNA linkers to be linear springs (stiffness $k_b$ and equilibrium length $l_b$) between cell and the substrate. In order to capture the mobile nature of these DNA linkers on the cell surface the DNA linkers on the cell surface are assumed to link the cell surface with the substrate location closest to the cell surface at the particular location. The adhesion force due to the DNA linkers is, therefore, given by

$$\mathbf{f}_a(s_0) = \begin{cases} -\rho_a(s_0) k_b \left(l/l_b - 1\right) \hat{\mathbf{e}}_y & \text{for } l \leq 2l_b \\ 0 & \text{for } l > 2l_b \end{cases} \tag{12}$$

where $l$ is the length of the adhesion bond between cell surface and substrate. Here we are assuming that there is no DNA linker bond of length greater than $2l_b$ and for $l < 2l_b$ the number of bonds are proportional to the DNA linker density. Please note that the locations on the cell surface where cytoskeletal stresses crosses the critical threshold value $\sigma_c$ the density of the DNA linkers $\rho_a$ is set to zero.

##### Pipette force

A portion of the dorsal surface of the cell is assumed to be in contact with the pipette through which the pulling force is applied normal to the contact surface (see the figure above). The contact between the pipette and cell surface is assumed to be perfect and there is no slipping between the two. In general the effect of pipette pull can be taken into account in two ways - force controlled or displacement controlled. Similar to the setup in the experiments in the model also we consider the pipette to be moving at a constant speed $v_p$ until the cell is completely detached from the substrate.

##### Non-dimensionalization and parameter values

We consider the radius of the unadhered cell $R_0$ to be the characteristic length of the system and we non-dimensionalize all the lengths in the system by $R_0$. Using cytoskeletal elastic and viscous contributions we can define characteristic timescale to be $\tau = \mu/k_e$. Since in the detachment process the viscous effects are smaller than the elasto-plastic ones we non-dimensionalize mechanical stress in the system by cortical stiffness $k_e$ and mechanical forces by $k_e t_c R_0$ where $t_c$ is the cortical thickness before adhesion of the cell. For all the results we have set non-dimensional values of parameters as $l_b = 0.02$, $k_b = 10.0$, $\zeta = 1.0$, $\sigma_c = 0.1$. The values of the rest of the parameters are specified in respective figure captions.

#### Simulation setup

This model of cell adhesion and detachment is numerically simulated for adhesion and detachment regimes. The strains in the cell surface during the cell detachment were calculated from the metric tensors (first fundamental form [14]) in the adhered and partially detached configurations. These tensors at a location $\mathbf{x}$ on the cell surface are defined as

$$g_0^{mn} = \frac{\partial \mathbf{x}_0^i}{\partial s_m} \frac{\partial \mathbf{x}_0^i}{\partial s_n} \tag{13}$$

and

$$g^{mn} = \frac{\partial \mathbf{x}_i}{\partial s_m} \frac{\partial \mathbf{x}_i}{\partial s_n} \tag{14}$$

where $s_m$ and $s_n$ are the parametrizations in two directions on the cell surface. From these tensors the principal extensions at location $\mathbf{x}$ can be calculated by diagnolization of the metric tensors by [15]

$$\det\left(g^{mn} - \epsilon^2 g_0^{mn}\right) = 0. \tag{15}$$

These relations linking cell surface geometry (in terms of $\mathbf{x}$) along with the constitutive model for the cytoskeleton (equations (7),(8),(9),(10) and (11)) and the external forces due to cell adhesion (given by equation (12)) and pipette force close the problem.

For computational implementation of the model the cell surface was discretized using triangular mesh as shown in the figure above.

##### Cell adhesion to the substrate

We assume that the cell adhesion takes place over a long time and during that period the cytoskeleton attains an equilibrium configuration. In other words, the adhesion of the cells is assumed to take place for a duration $\gg \tau$. Therefore, during this time the nature of the cytoskeleton is considered to be dominated by its viscous flow. At the start of the simulations of cell adhesion we assume a homogeneous distribution of actin, myosin and DNA binders on the cell surface except a very small inhomogeneity (to break the symmetry) at the center of the contact zone. Contractile forces generated by the myosin result in the flow of the cytoskeleton along the cell surface which also results in the flow of myosin and adhesion molecules.

##### Cell detachment from the substrate

Once the cell is adhered to the substrate and equilibrium is reached we apply constant rate of displacement at some nodes of the dorsal surface of the cell. This rate of displacement is kept very small to ensure the quasi-static nature of the system holds true. However during this time the dominant contribution of the cytoskeleton is considered to be from its elasto-plasticity. In other words during this stage we do not consider any viscous flow of the cytoskeleton relative to the cell surface. This assumption can be justified by the fact that in experiments the cell detachment takes place over a very short period of time.

[5] Ron Milo and Rob Phillips. *Cell biology by the numbers.* Garland Science, 2015.

[14] Erwin Kreyszig. *Differential geometry.* Courier Corporation, 2013.

[15] Alexander Farutin, Thierry Biben, and Chaouqi Misbah. 3d numerical simulations of vesicle and inextensible capsule dynamics. *Journal of Computational Physics*, 275:539–568, 2014.

### Supplementary results

#### Actin cortex thickness correlates with adhesion toughness

We asked if previously unexplained disparity in adhesion toughness between different cell types can be explained by presence of a more prominent cortex which intuitively may be able to store and dissipate more energy. We therefore compared two cell types, A431D and S180. Both cell types are endogenously deprived of any cadherin expression. We created stable cell lines where A431D has a significantly higher expression of E-cad on its plasma membrane (48 au) as compared to S180 (14 au) (**Supplementary Figure 7a**). By contrast A431D displayed a much lower amount of cortical actin (17 au) (phalloidin staining, see **Material and Methods)**, as compared to S180 (93 au) (**Supplementary Figure 7b**). Contrary to our present understanding where cadherin expression is often co-related with adhesion strength, it was observed that the junction cohesion was about twice stronger for S180 than for A431D (separation force (115nN *vs* 32nN), strain to failure (46% *vs* 32%), and adhesion toughness (28 x $10^7$ kT *vs* 11 x $10^7$ kT) (**Supplementary Figure 7d-f**). These data strongly confirm that cortical actin network was a much stronger predictive factor of adhesion strength of cells than cadherin expression level.

#### EGFR signalling and cortex modulation

Previous studies have showed involvement of EGFR signalling at adherens junction. We performed a pan-tyrosine kinase wide screen to evaluate activation of EGFR during E-cad mediated cell-cell adhesion. Our results confirmed robust activation of EGFR after adherens junction formation using both immunoblot and western blotting for pEGFR (**Supplementary figure 8a-c**). This response, however, did not occur on cell aggregation mediated through DNA-cad (**Supplementary figure 8d**). Further confirming that DNA-cad lacked the ability to modulate cell signalling in a way analogous to E-cad. We reasoned that the main driver that controls the cohesion could be the cytoskeleton properties independently of the binder recruitment. We tested this hypothesis by testing $DNA_{45}$-cad cells activated with a burst of soluble EGF (100 ng/ml), a known activator of Rho GTPases Rac1 and Cdc42 (*22*), to compensate the deficient activation of this pathway that we identified in DNA-cad. EGF treatment did not change significantly the junction morphology (**Supplementary figure 9a**), binder distribution at the junction ((**Supplementary figure 9b**), amount of binder at the junction (**Supplementary figure 9c**), or the junction size (**Supplementary figure 9d**). Interestingly, EGF treatment rescued $F_s$ (P=0.89), and U (P=0.35) to their control values (E-cad) (**Supplementary figure 10a, b, d),** and increased $\varepsilon$ to a significantly higher level as compared to E-cad control (P=0.001) (**Supplementary figure 10 c**). Furthermore, it was seen that while EGF treatment alone lead to a two-fold increase in the cortical tension of cells (**Supplementary figure 10e**), simultaneous treatment of EGF and Nocodazole 10 µM showed a phenotype wherein EGF could buffer the nocodazole mediated rise in cortical tension and rigidity of cells. This was in line with our data on ability of N-WASP to prevent uncontrolled rise in cell rigidity even upon treatment with high concentration of nocodazole. This could also indicate that EGFR could be one of the pathways that the cell employs to modulate the cortex while signalling from adherens junction.

**Supplementary table 1**: Comparison between different features of E cadherin and engineered DNA-cadherin system

| Criteria | E cadherin | DNA-cadherin |
|---|---|---|
| Energy per binder | Fixed | Depends on the length and sequence of DNA |
| Formation kinetics (t) | 5 mins | 6 mins (ns) |
| Junction size | Tension dependent | Tension dependent |
| Cadherin localization | Tension dependent | Tension dependent |
| Mechanosensitive recruitment | Sensitive | Insensitive |
| Cortical tension change post junction formation | Yes (1.6 ± 0.8) | No (0.92±0.57) |

### Supplementary Figures

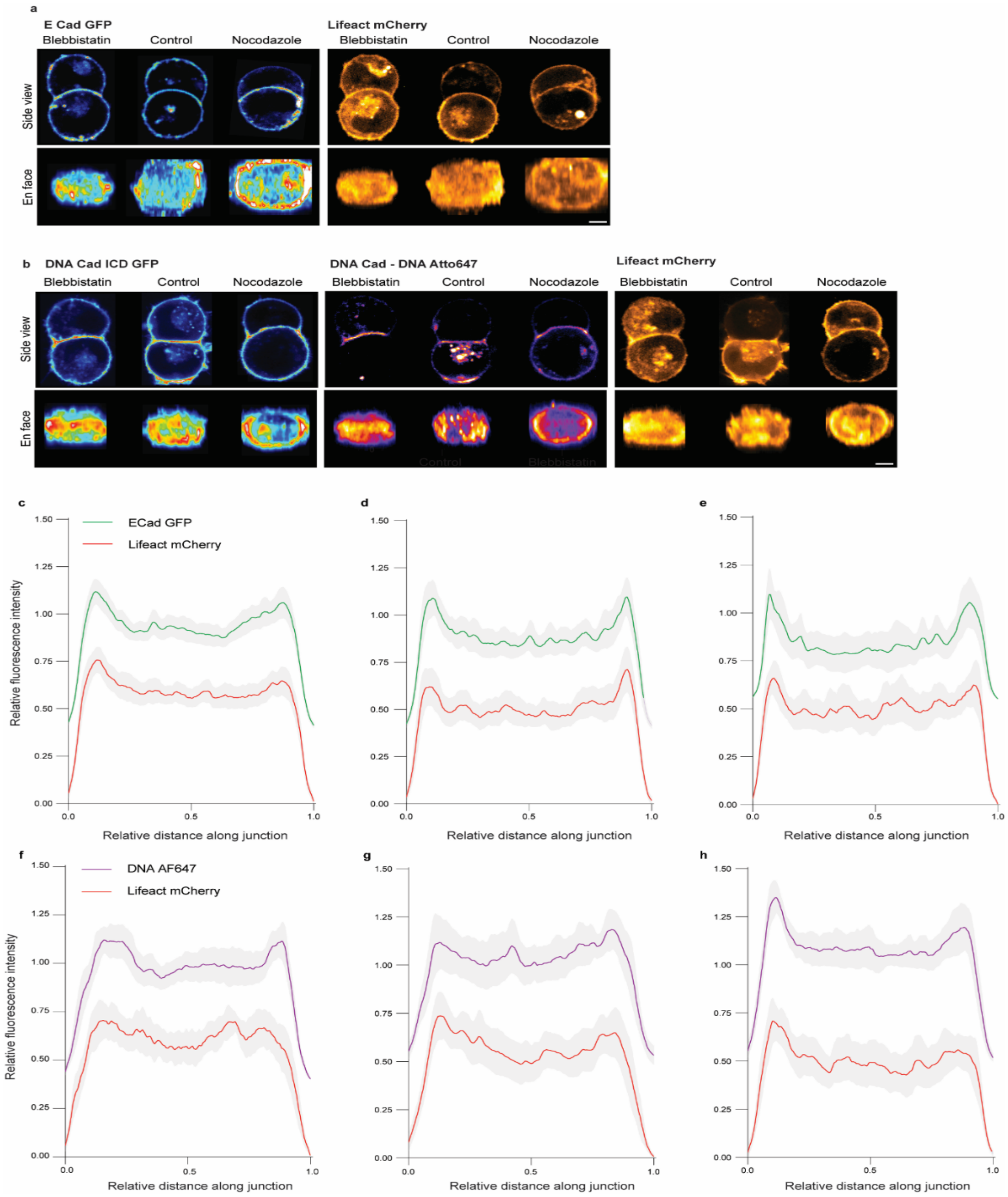


**Supplementary figure 1**: - **The dependence of cadherin and cortical actin distribution at cell-cell contact with cortical tension**. (a) E-cad GFP and Lifeact mCherry distribution at the *en-face* and side views of the E-cad GFP cell-cell contact pA-Blebbistatin, control and Nocodazole (b) DNA AF 647 and Lifeact mCherry distribution at the *en-face* and side views of the DNA-cad cell-cell contact under pA-Blebbistatin, control and Nocodazole (c-e) Average E-cad GFP and Lifeact mCherry fluorescence intensity distribution along the length of cell-cell contact under blebbistatin (c), control (d) and nocodazole (e). (f-h) Average DNA AF647 and Lifeact mCherry fluorescence intensity distribution along the contact length under blebbistatin (f), control (g) and nocodazole (h).

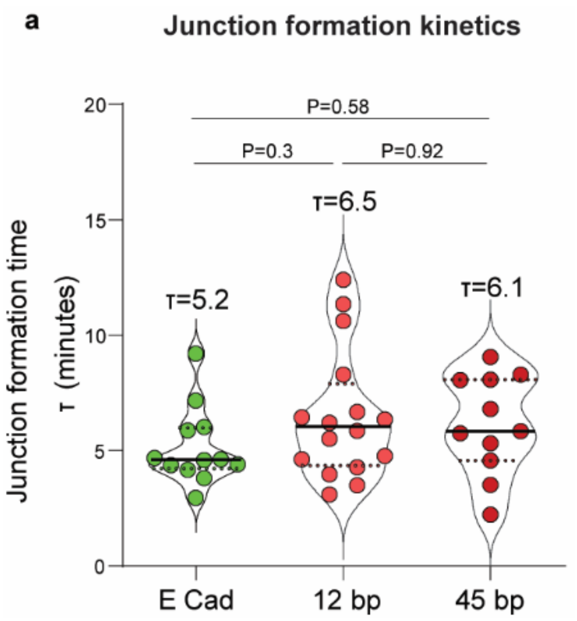


**Supplementary figure 2**: **Junction Expansion kinetics of E-cad and DNA-cad cell-cell doublets.** Junction formation time (τ) of E-cad cell-cell doublets and $DNA_{12}$-cad and $DNA_{45}$-cad cell-cell doublets.

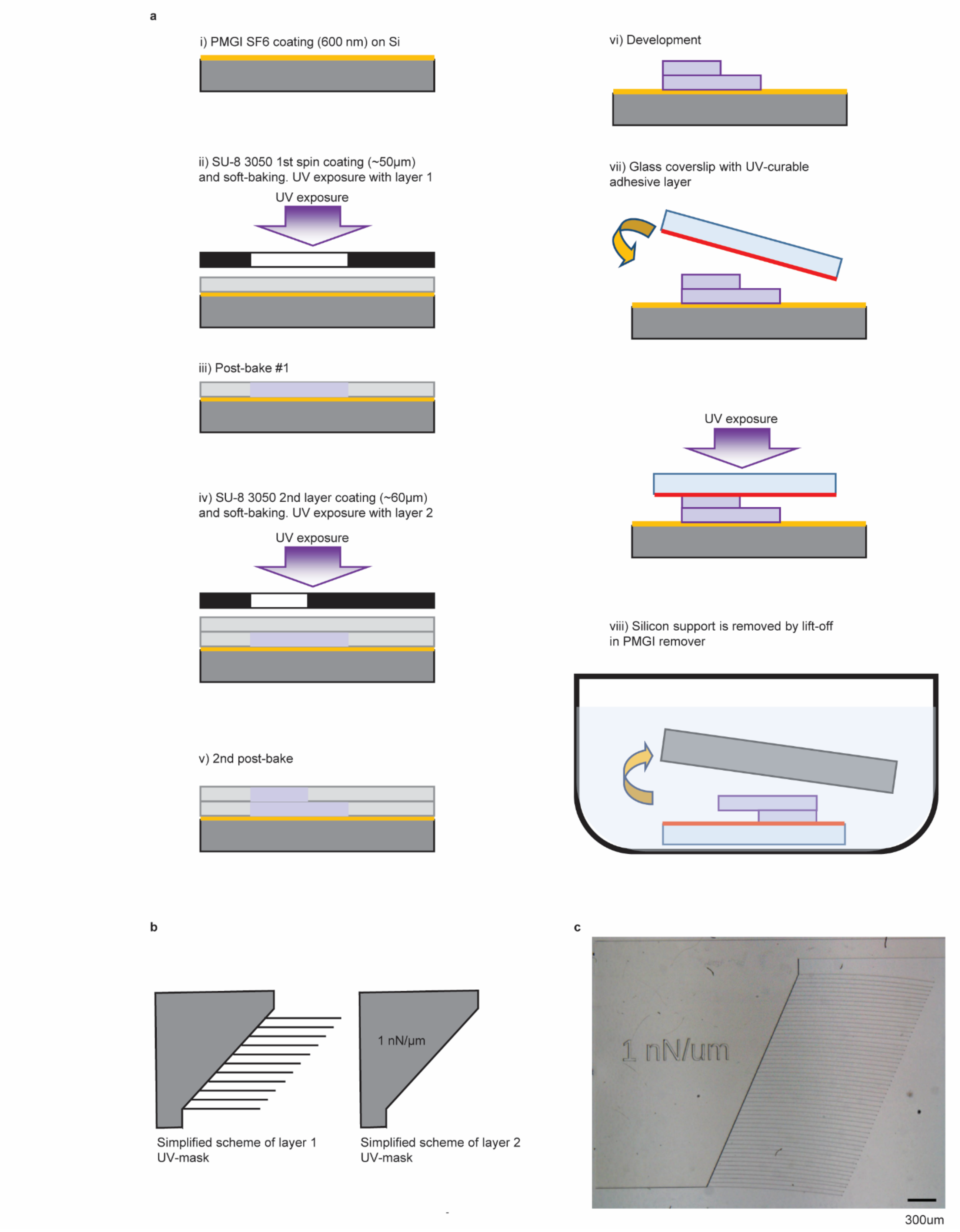


**Supplementary figure 3**: **Fabrication of SU-8 cantilevers using aligned lithography.** (a) Schematic depicting steps during the fabrication of cantilevers using two-layer aligned lithography and transfer to their glass coverslip. (b) Simplified scheme of UV-mask for the two layers of cantilever device. (c) Brightfield micrograph of the fabricated cantilever device. Scale bar 300 µm.

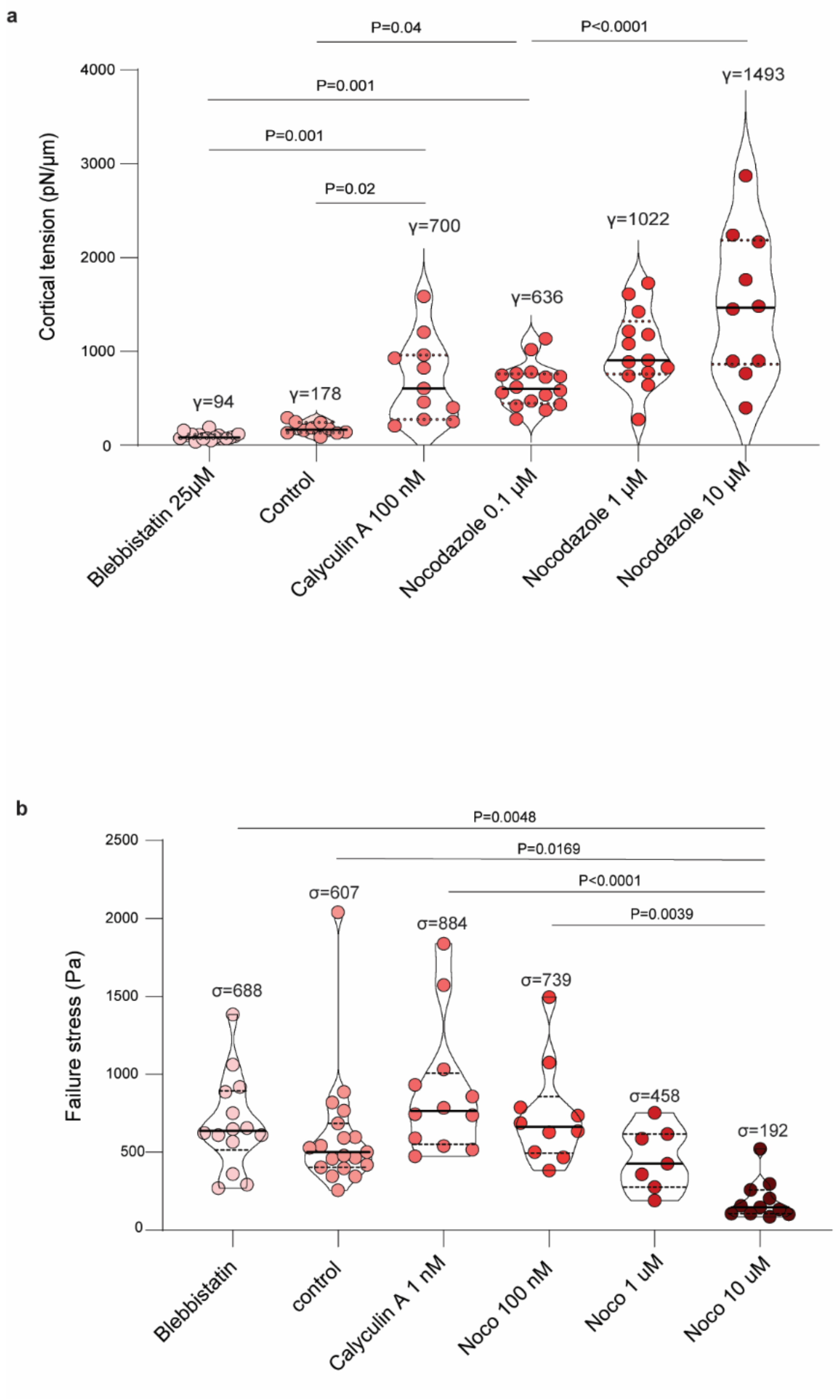


**Supplementary figure 4**: **Dependence of the cortical tension and fracture stress under small compound treatment.** (a) Quantification of cortical tension for DNA-cad doublets under different treatment conditions to modulate contractility. (b) Quantification of stress at fracture (Separation Force/ contact area) for DNA-cad doublets under different treatment conditions to modulate contractility.

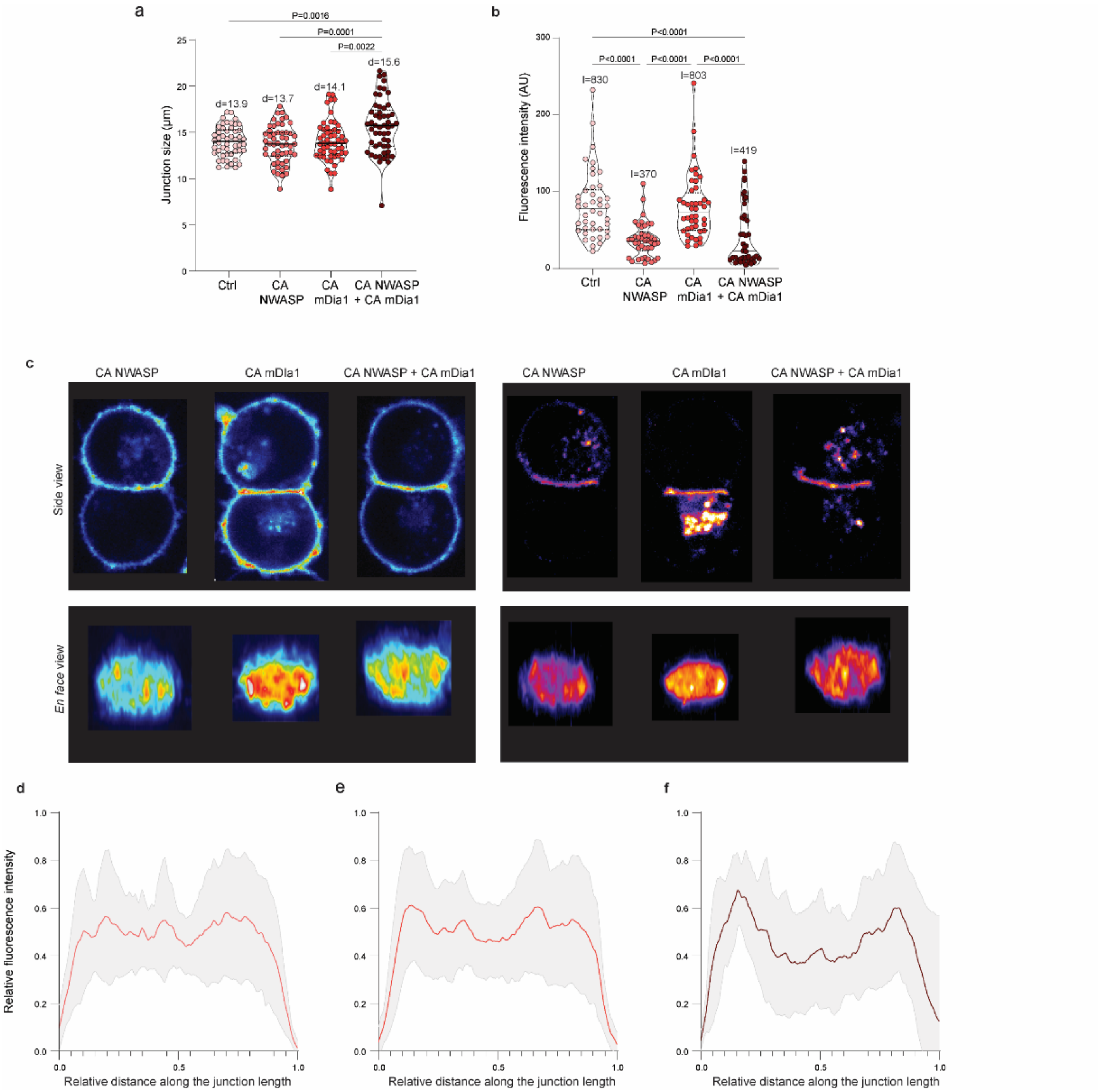


**Supplementary figure 5: Influence of actin nucleators on DNA-cad adhesion characteristics**. (a) Junction size for DNA-cad doublets with CA N-WASP, CA mDia1 and both CA N-WASP and CA mDia1 in comparison with control. (b) Fluorescence intensity of DNA AF647 for DNA doublets expressing CA N-WASP, CA mDia1 and both CA N-WASP and CA mDia1 in comparison with control. (c) Side and *en face* view of DNA-cad cell doublets (Cad ICD GFP, and DNA – Alexa fluor 647) expressing CA N-WASP, CA mDia1 and both CA N-WASP and CA mDia1. (d), (e) and (f) Average DNA AF647 fluorescence intensity distribution along the length of cell-cell contact for DNA doublets expressing CA N-WASP, CA mDia1 and both CA N-WASP and CA mDia1 respectively.

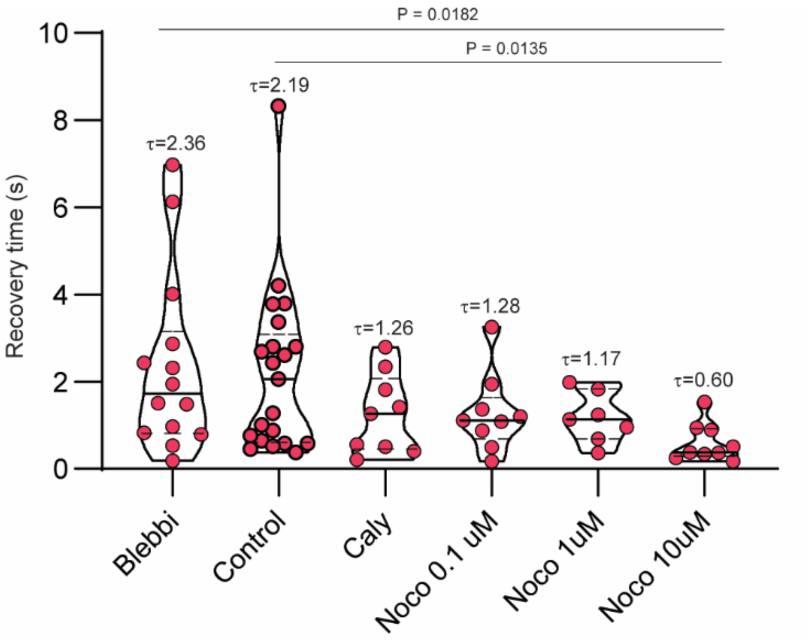


**Supplementary figure 6: Influence of different drug treatments to perturb contractility on cell shape recovery time (τ in secs) after fracture.**

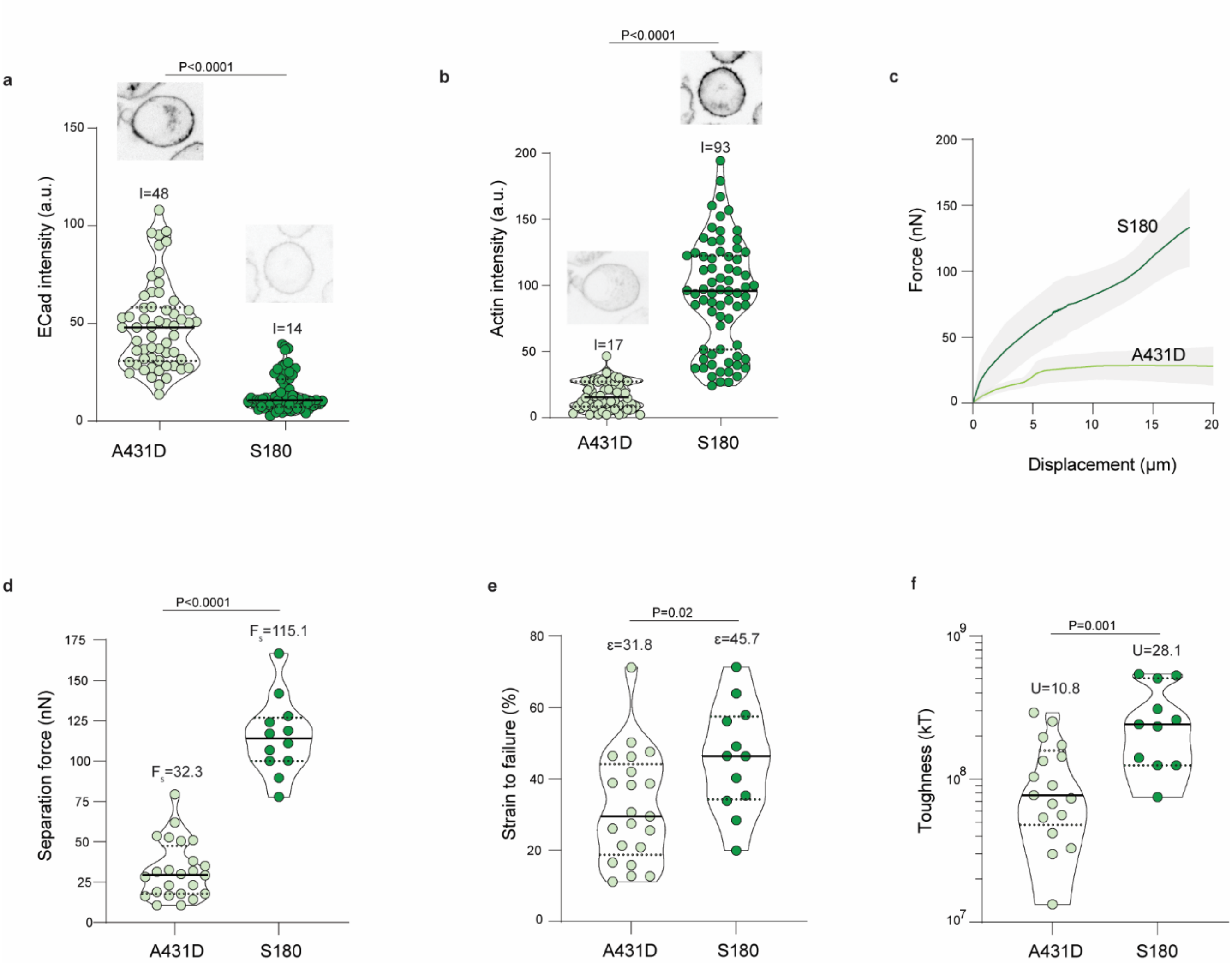


**Supplementary figure 7: Cortical actin as the potent regulator of cell adhesion toughness**. (a-b) Total E cadherin (a) and actin (b) intensities in A431D E-cad GFP and S180 cells. Cells with higher cortical actin but lower E cadherin intensity (s180) shows higher force-displacement curve area (c), higher separation force (d), higher strain to failure rate (e) and higher toughness (f) as opposed of cells with lower cortical actin and higher E cadherin expression (A431D E-cad GFP).

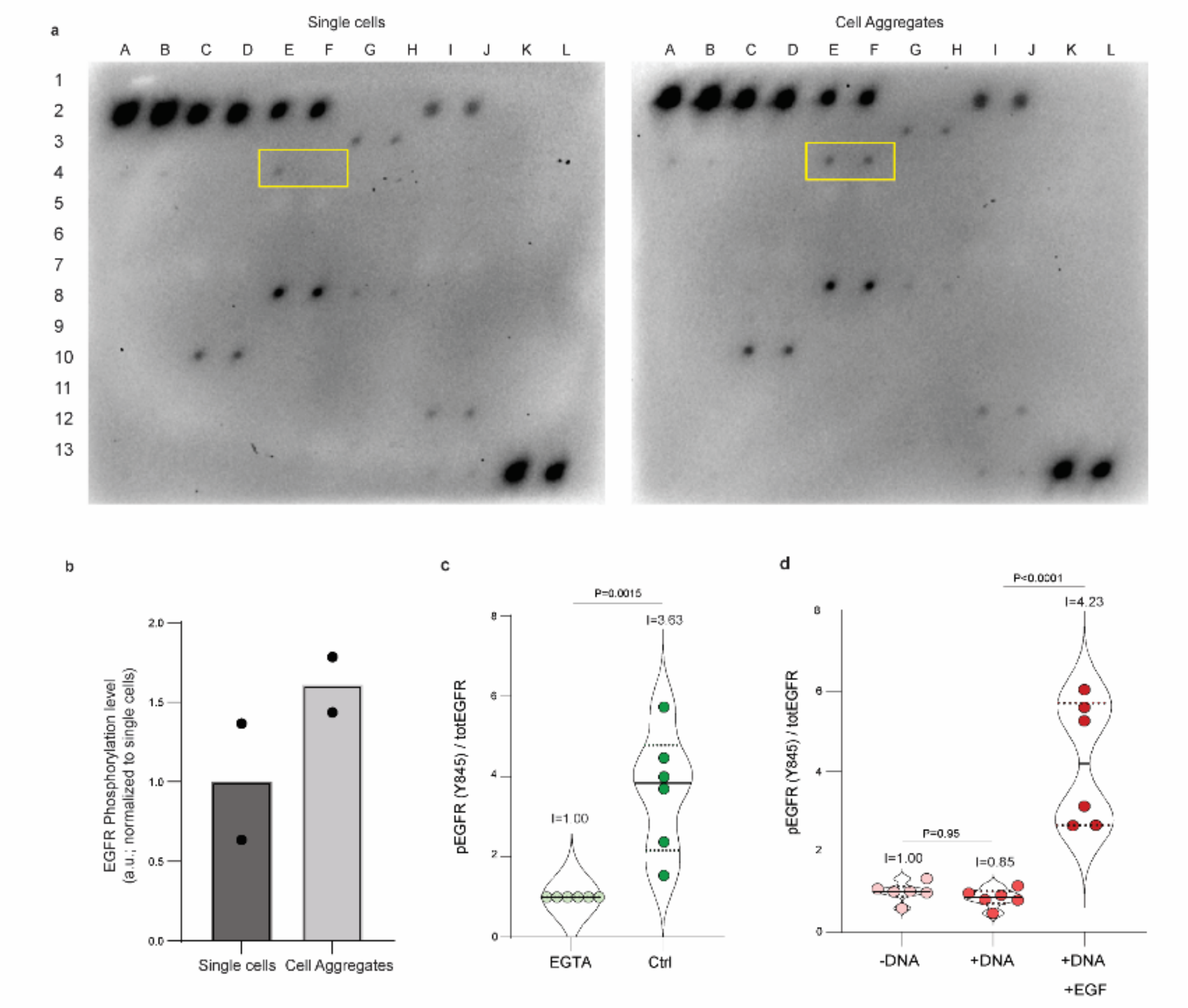


e

| | A | B | C | D | E | F | G | H | I | J | K | L |
|---|---|---|---|---|---|---|---|---|---|---|---|---|
| 1 | POS1 | POS1 | POS2 | POS2 | POS3 | POS3 | ABL1 | ABL1 | ACK1 | ACK1 | ALK | ALK |
| 2 | NEG | NEG | NEG | NEG | Axl | Axl | Blk | Blk | BMX | BMX | Btk | Btk |
| 3 | Csk | Csk | Dtk | Dtk | EGFR | EGFR | EphA1 | EphA1 | EphA2 | EphA2 | EphA3 | EphA3 |
| 4 | EphA4 | EphA4 | EphA5 | EphA5 | EphA6 | EphA6 | EphA7 | EphA7 | EphA8 | EphA8 | EphB1 | EphB1 |
| 5 | EphB2 | EphB2 | EphB3 | EphB3 | EphB4 | EphB4 | EphB6 | EphB6 | EerB2 | EerB2 | EerB3 | EerB3 |
| 6 | EerB4 | EerB4 | FAK | FAK | FER | FER | FGFR1 | FGFR1 | FGFR2 | FGFR2 | FGFR2 (α) | FGFR2 (α) |
| 7 | Fgr | Fgr | FRK | FRK | Fyn | Fyn | Hck | Hck | HGFR | HGFR | IGF-IR | IGF-IR |
| 8 | Insulin R | Insulin R | Itk | Itk | JAK1 | JAK1 | JAK2 | JAK2 | JAK3 | JAK3 | LCK | LCK |
| 9 | LTK | LTK | Lyn | Lyn | MATK | MATK | M-CSFR | M-CSFR | MUSK | MUSK | NGFR | NGFR |
| 10 | PDGF R-α | PDGF R-α | PDGF R-β | PDGF R-β | PYK2 | PYK2 | RET | RET | ROR1 | ROR1 | ROR2 | ROR2 |
| 11 | ROS | ROS | RYK | RYK | SCFR | SCFR | SRMS | SRMS | SYK | SYK | Tec | Tec |
| 12 | Tie-1 | Tie-1 | Tie-2 | Tie-2 | TNK1 | TNK1 | TRKB | TRKB | TXK | TXK | NEG | NEG |
| 13 | Tyk2 | Tyk2 | TYRO10 | TYRO10 | VEGFR2 | VEGFR2 | VEGFR3 | VEGFR3 | ZAP70 | ZAP70 | POS4 | POS4 |

**Supplementary figure 8: Immunoblot-based screening for receptor tyrosine kinase phosphorylation using antibody array membrane.** (a) Immunoblot array for single cells, and post-cell aggregation. The array positions for EGFR are highlighted using the yellow boxes. (b) Quantification of pTyr intensity for EGFR. (c) Reduction in pEGFR through EGTA blockage of homophilic trans E-cad interaction (d) Quantification of pEGFR with and without DNA and also with DNA and EGF. (e) Position map of RTK immunoblot array used in the screening experiment.

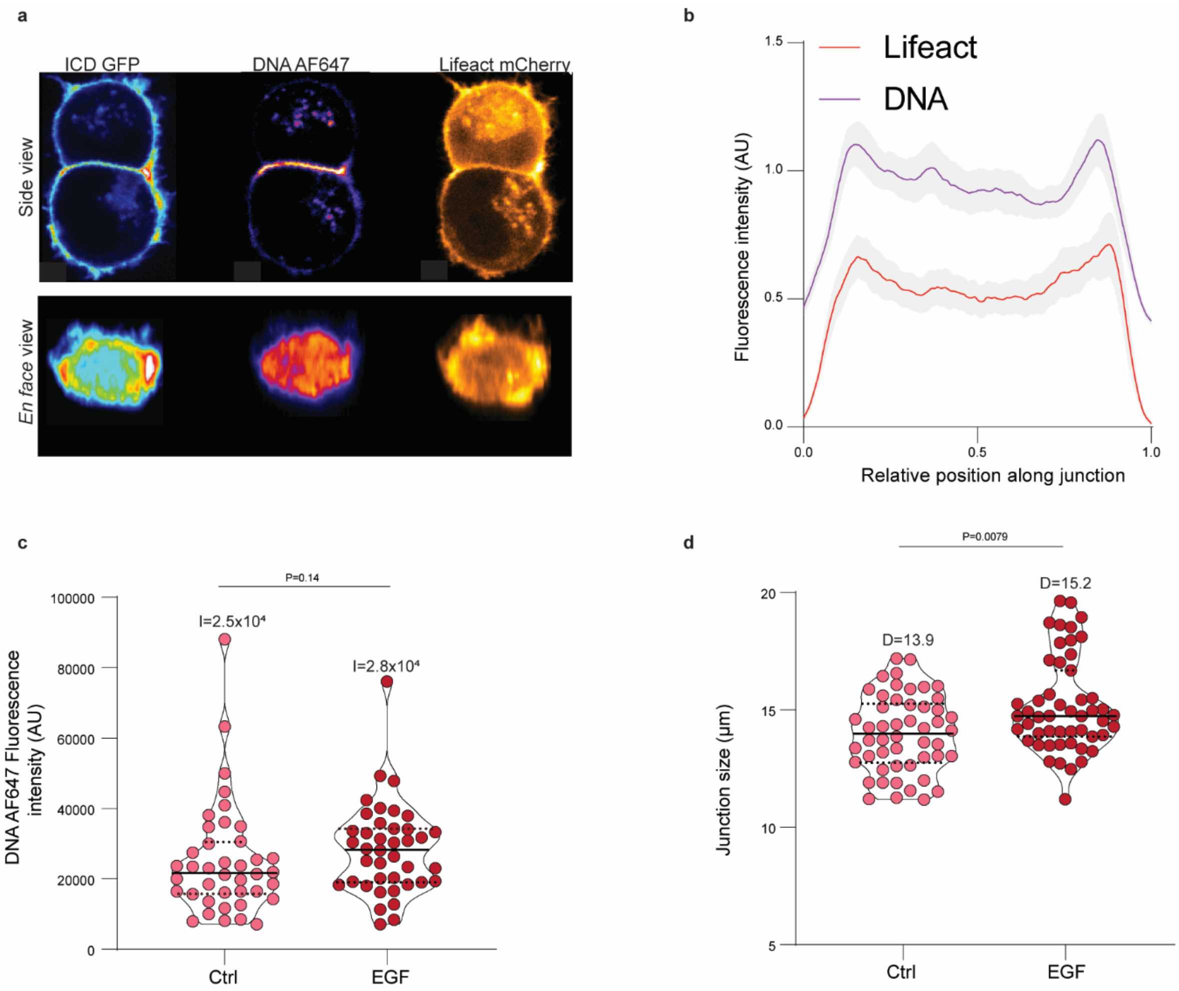


**Supplementary figure 9: Effect of EGFR activation using soluble EGF treatment on junction organization in DNA-cad doublets**. (a) Fluorescence intensity of DNA AF647 and LifeAct mCherry (b) Average distribution of AF647 DNA and LifeAct mCherry along the length of cell-cell junctions (c) Quantification of AF647 DNA fluorescence intensity upon addition of EGF (d) Quantification of junction size upon EGF treatment.

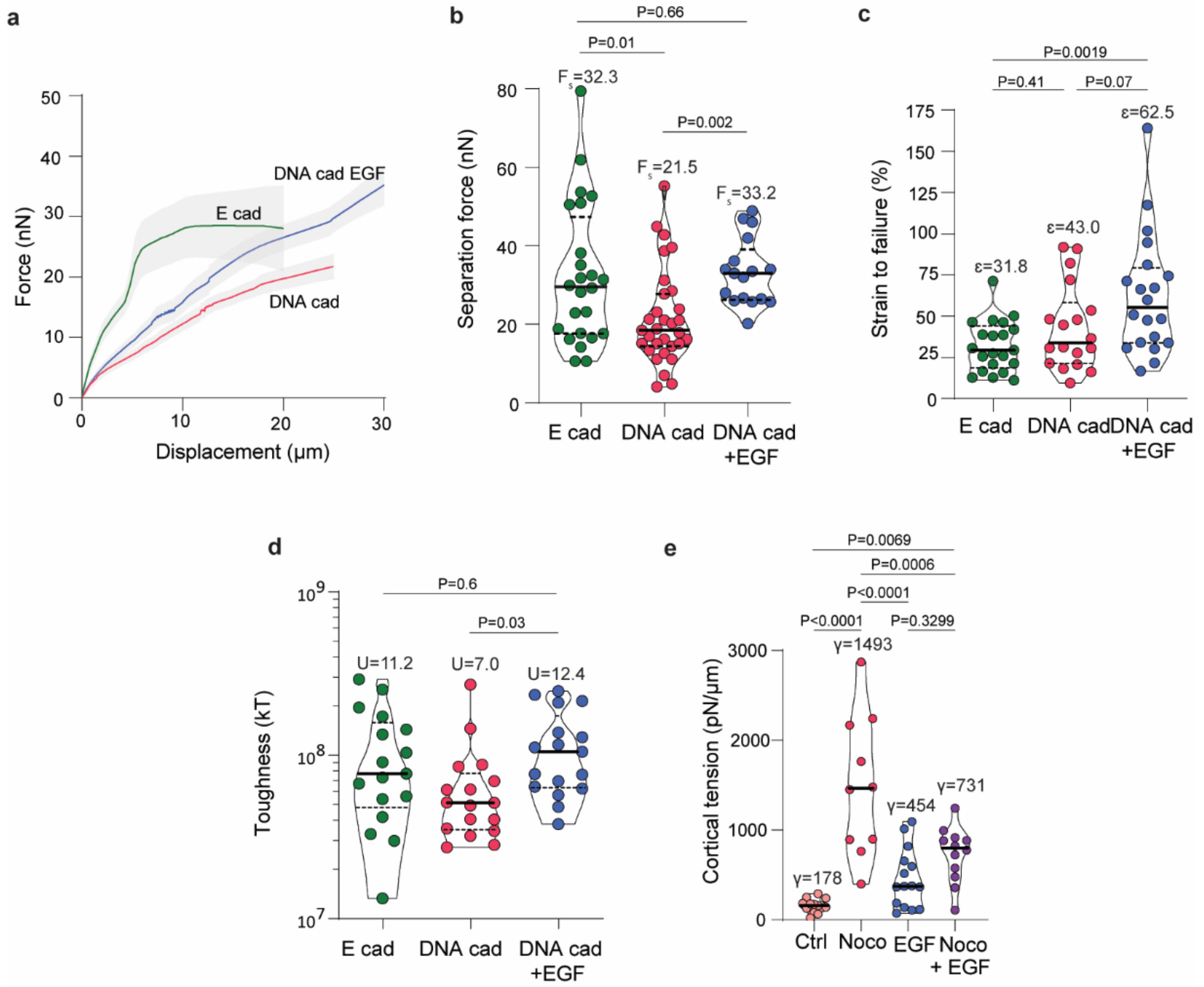


**Supplementary figure 10: Effect of EGFR signaling on junction separation force, cell ductility and toughness.** Partial rescue of the force-displacement area curve (a) and separation force (b) upon EGF addition to DNA-cad. (c) Higher strain to failure rate upon EGF addition to DNA-cad. Partial rescue of toughness (d) and increased cortical tension (e) upon Nocodazole treatment through addition of EGF to DNA-cad.

a

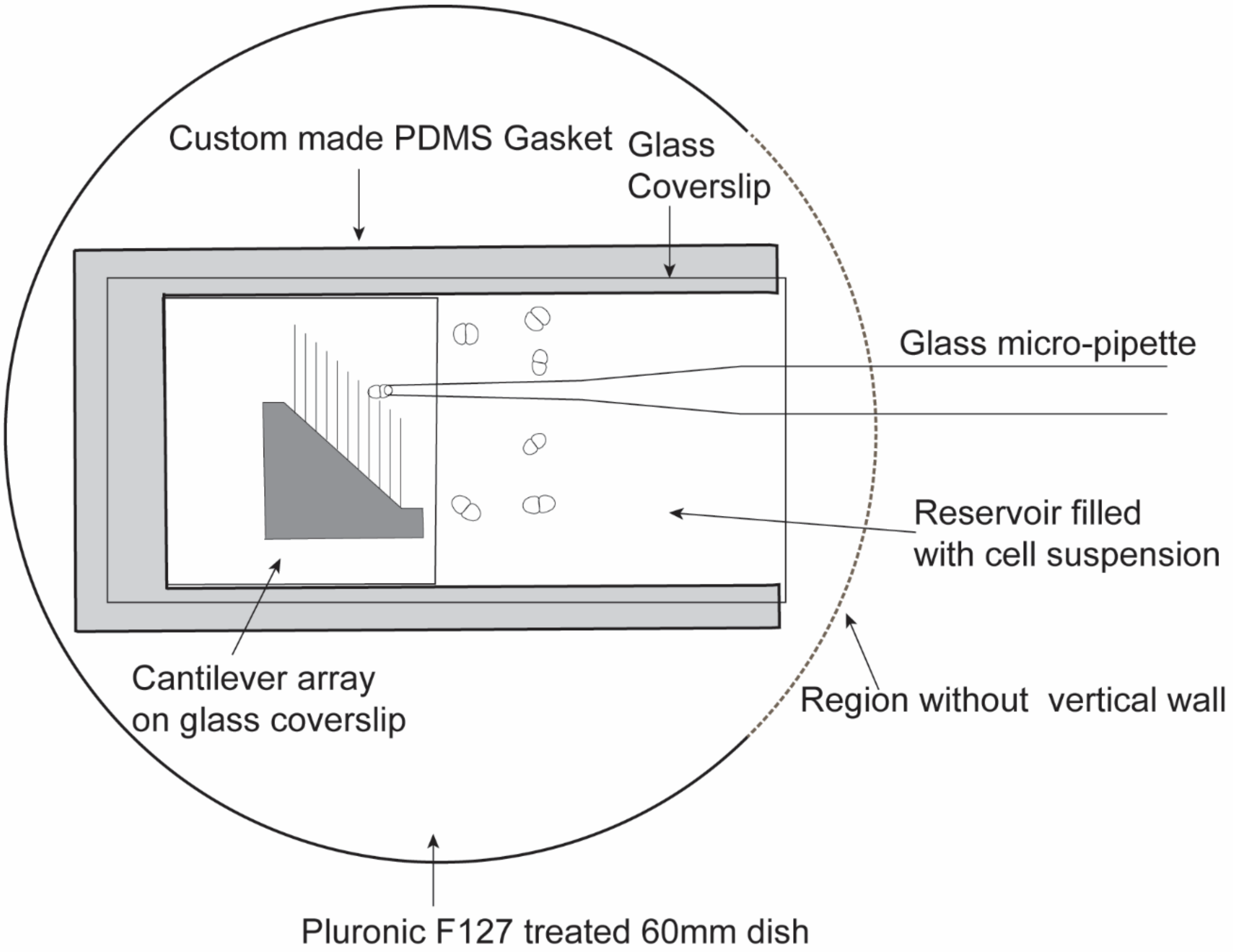


b

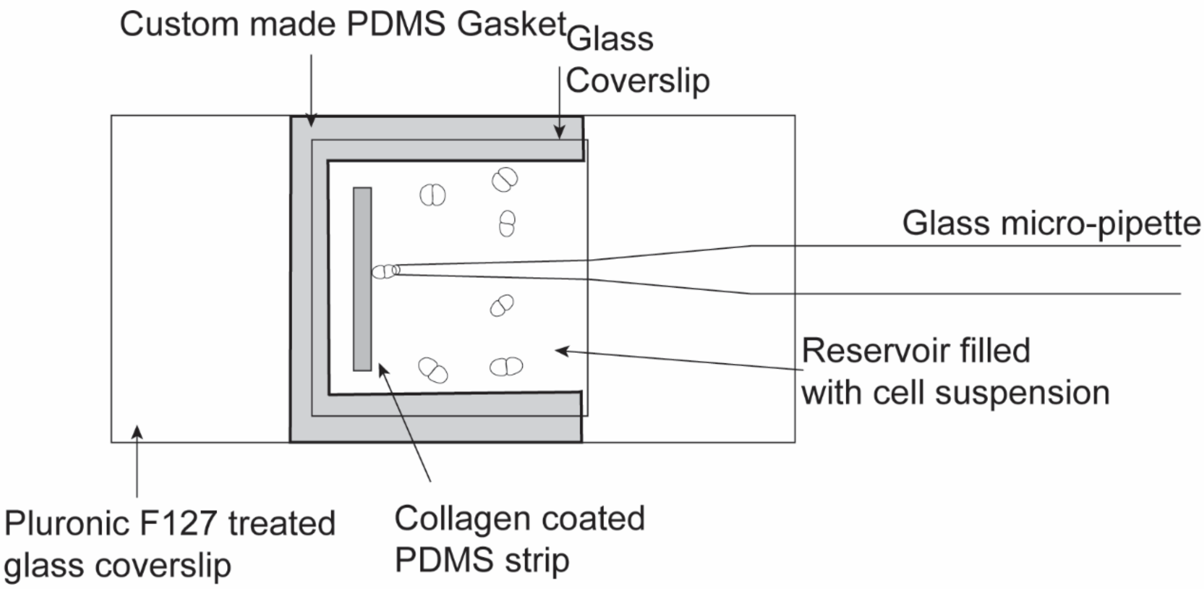


**Supplementary figure 11: Schematic of experimental setup for studying cell-cell adhesion mechanics.** (a) Experimental setup for studying cell-cell adhesion mechanics using cantilever array. (b) Experimental setup for fluorescence imaging of cell-cell adhesion during fracture.

### Supplementary Video

**Supplementary Video 1.** Cell-cell doublets formed by DNA-cad expressing cells undergoing separation on cantilever setup. Scale bar 10 µm.

**Supplementary Video 2.** Cell-cell doublets formed by DNA-cad expressing cells treated with 100 nM Nocodazole undergoing separation on cantilever setup. Scale bar 10 µm.

**Supplementary Video 3.** Cell-cell doublets formed by DNA-cad expressing cells treated with 10 µM Nocodazole undergoing separation on cantilever setup. Scale bar 10 µm.

**Supplementary Video 4.** Maximum intensity projections of cell-cell doublets formed by DNA-cad expressing cells undergoing separation. Cad ICD GFP (Royal LUT); DNA Atto647 (Fire LUT). Scale bar 5 µm.

**Supplementary Video 5.** Maximum intensity projections of cell-cell doublets formed by DNA-cad expressing cells treated with 10 µM nocodazole undergoing separation. Cad ICD GFP (Royal LUT); DNA Atto647 (Fire LUT). Scale bar 5 µm.

**Supplementary Video 6.** Maximum intensity projections of cell-cell doublets formed by DNA-cad expressing cells treated with 10 µM Nocodazole undergoing separation. Cad ICD GFP (Royal LUT); SPY-FastAct650 (Orange hot LUT). Scale bar 5 µm.

**Supplementary Video 7.** Cell-cell doublets formed by DNA-cad-CA-N-WASP expressing cells undergoing separation on cantilever setup. Scale bar 10 µm.

**Supplementary Video 8.** Cell-cell doublets formed by DNA-cad + CA-mDia1 expressing cells undergoing separation on cantilever setup. Scale bar 10 µm.

**Supplementary Video 9.** Cell-cell doublets formed by DNA-cad-CA-N-WASP + CA-mDia1 expressing cells undergoing separation on cantilever setup. Scale bar 10 µm.

**Supplementary Video 10.** Maximum intensity projections of cell-cell hetero-doublets formed between DNA-cad and DNA-cad-CA-N-WASP expressing cells treated with 10 µM Nocodazole undergoing separation. Cad ICD GFP (Royal LUT); DNA Atto647 (Fire LUT). Scale bar 5 µm.

**Supplementary Video 11.** DNA-cad expressing cell treated with 10 µM nocodazole undergoing micropipette aspiration. Scale bar 5 µm.

**Supplementary Video 12.** DNA-cad-CA-N-WASP expressing cell treated with 10 µM nocodazole undergoing micropipette aspiration. Scale bar 5 µm.

**Supplementary Video 13.** Total stress mapped in the en face and side view of cells with different contractilities undergoing de-adhesion from simulation experiments described in Figure 6.

**Supplementary Video 14.** Results from simulation experiments depicting local strain for cells having different elasto-plastic characteristics as in Figure 6.
